# DNA-Origami-Based Fluorescence Brightness Standards for Convenient and Fast Protein Counting in Live Cells

**DOI:** 10.1101/2020.09.20.305359

**Authors:** Nathan D. Williams, Ane Landajuela, Ravi Kiran Kasula, Wenjiao Zhou, John T. Powell, Zhiqun Xi, Farren J. Isaacs, Julien Berro, Derek Toomre, Erdem Karatekin, Chenxiang Lin

**Affiliations:** Department of Cell Biology, Yale University School of Medicine, New Haven, CT 06520, USA; Nanobiology Institute, Yale University, West Haven CT 06516, USA; Department of Cellular and Molecular Physiology, Yale University School of Medicine, New Haven, CT 06520, USA; Department of Molecular, Cellular and Developmental Biology, Yale University, New Haven, CT 06520, USA; Department of Biomedical Engineering, Yale University, New Haven, CT 06520, USA; Systems Biology Institute, Yale University, West Haven, CT 06516, USA; Department of Molecular Biophysics and Biochemistry, New Haven, CT 06520, USA; Université de Paris, SPPIN – Saints-Pères Paris Institute for the Neurosciences, Centre National de la Recherche Scientifique (CNRS), F-75006 Paris, France

## Abstract

Fluorescence microscopy has been one of the most discovery-rich methods in biology. In the digital age, the discipline is becoming increasingly quantitative. Virtually all biological laboratories have access to fluorescence microscopes, but abilities to quantify biomolecule copy numbers are limited by the complexity and sophistication associated with current quantification methods. Here, we present DNA-origami-based fluorescence brightness standards for counting 5–300 copies of proteins in mammalian and bacterial cells, tagged with fluorescent proteins or organic dyes. Compared to conventional quantification techniques, our brightness standards are robust, straightforward to use, and compatible with nearly all fluorescence imaging applications, thereby providing a practical and versatile tool to quantify biomolecules *via* fluorescence microscopy.

---

Fluorescence microscopy is widely used in biology and biomedicine as a tool to visualize cell morphology and dynamics, in particular the biomolecules (e.g., DNA, RNA, protein) of interest that are labeled with a fluorophore. In modern life science studies, it is increasingly important to not only know the identities and locations of biomolecules, but also to measure their stoichiometry. However, methods to count biomolecules from fluorescence images usually lack versatility (e.g., they are limited to certain imaging modalities or dynamic range) and/or require sophisticated analysis. Stepwise photobleaching, while precise under finely tuned conditions, is limited to small (≲10) numbers of molecules.^1–6^ Quantitative immunoblot-calibrated internal fluorescence standards can quantify molecules of a wide range of copy numbers (from tens to thousands)^7–10^ but may suffer batch-to-batch variations that necessitate repeated, laborious calibrations. Super-resolution methods offer an increase in resolution and precision, but often at the expense of ease of use (i.e., specialized sample preparation and/or microscope) and speed (longer data acquisition and processing time).^11–14^ Therefore, there is a pressing need for a fast, universal technique that can be conveniently integrated into a wide array of existing imaging workflows to count biomolecules. A customizable external standard would fit this description: it would require little to no additional cloning, could be programmed to count virtually any fluorophore, and sustain long-term storage before being imaged in physiologically relevant conditions. As a precise, programmable, and robust self-assembly method, DNA origami^15–18^ is ideal for the creation of such a standard. DNA origami has been used to engineer standards for fluorescence microscopy, but existing quantification methods either rely on super-resolution techniques or only demonstrate applications for counting organic dyes.^12,19^ Here, we present DNA-origami-based brightness standards that incorporate 5–200 organic dyes and fluorescent proteins and use them for quantitative fluorescence microscopy on bacterial and mammalian cells. Our fluorescent standards offer a fast and precise solution to counting biomolecules in different cell types, with commonly used microscopes and fluorophores, including fluorescent proteins expressed in cells.

The design of our DNA-origami structures for generating brightness standards is based on the well-documented 6 helix-bundle (6hb) nanotube, which is ∼7 nm in diameter and ∼407 nm long^15,20–22^ (**Figure 1a** & **S1**–**S4**). Previously, 6hb and other rod-shaped DNA-origami structures have been used to construct fluorescence markers and barcodes for bioimaging,^12,22–26^ owing to their robust assembly (up to 90% yield) as well as their thermal (melting temperature ≈ 60°C)^27^ and mechanical stability (persistence length>1 µm).^28^ In these applications, single-stranded extensions, termed handles, are placed at designated positions on the surface of DNA nanotubes with nanometer precision to host a myriad of fluorophores. In this work, handles are precisely positioned at 42 bp or ∼14 nm apart on each helix to maximize labeling density while minimizing self-quenching. As such, each 6hb structure can accommodate up to 100 copies of fluorophores of interest (e.g., monomeric enhanced green fluorescent protein, mEGFP) in the center, with 12 additional handles at each end reserved for other fluorophores with distinct emission spectra (e.g., Alexa Fluor 647) to aid focusing and quality control. Additionally, 4 handles are evenly spaced on one of the helices to display biotin labels for surface attachment. We designed five versions of 6hb structures to carry 5, 25, 50, 70, and 100 mEGFPs (**Figure 1a**), which we prepared from a 7308-nt long circular ssDNA and 5 different pools of synthetic oligonucleotides following well-established DNA-origami assembly and purification protocols.^16,29–32^

**Figure 1.**
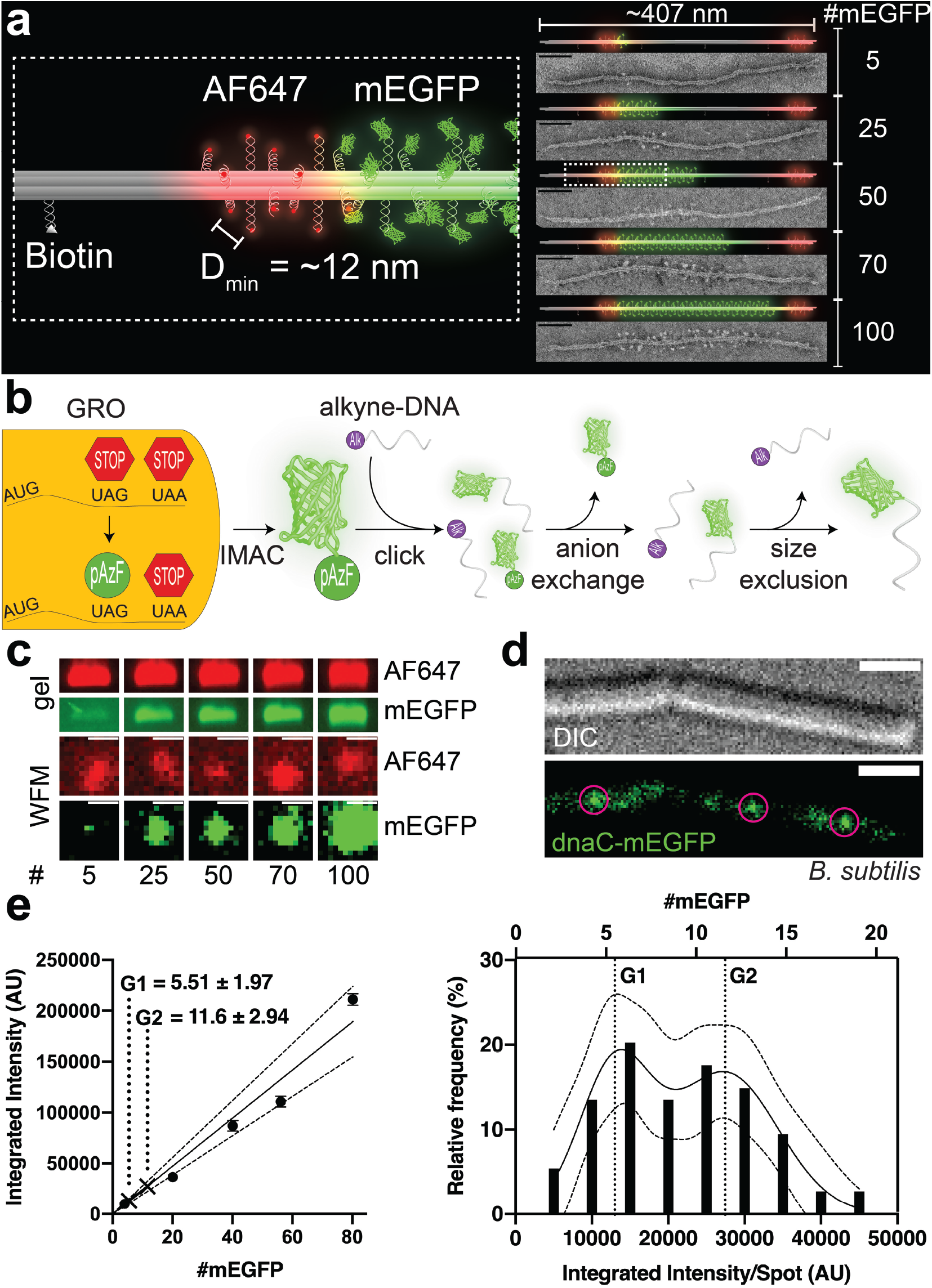
A DNA-origami-based mEGFP brightness standard. (a) 3D models and TEM micrographs of monomeric DNA 6hb structures labeled with 5–100 copies of mEGFP (green) in the main body, 12 copies of Alexa Fluor 647 (red) at each end, and 4 biotin molecules along one side. The minimum spacing of the fluorophores is ∼12 nm. Scale bars: 50 nm. (b) Generation of mEGFP-DNA conjugate. mEGFP-pAzF is expressed and purified from a GRO, in which the antisense TAG codon has been reassigned to encode pAzF, an azide-modified Phe. mEGFP(pAzF) was purified *via* immobilized metal affinity chromatography (IMAC), then reacted with alkyne-labeled DNA. Two subsequent purification steps removed unreacted proteins and DNA. (c) Gel electrophoresis (top) and widefield microscopy images (WFM, bottom) of mEGFP standards. Images are set to the same brightness scale (no saturated pixels in the original images). Scale bars: 2 µm. (d) Differential interference contrast (DIC, top) and WFM (bottom) images of *B. subtilis* (stain NW001) expressing dnaC-mEGFP. Circles indicate puncta picked for quantification. Scale bars: 2 µm. (e) Quantifying dnaC-mEGFP. Left: a calibration curve with intensities of DNA-origami standards and interpolated protein counts (mean±SEM) from dnaC-mEGFP puncta. Dotted lines denote 95% confidence interval. Right: frequency distribution and sum-of-two-Gaussians fit of dnaC-mEGFP puncta.

To generate DNA-conjugated mEGFP, we used a Genomically Recoded Organism (GRO) to express mEGFP with a single azide-bearing nonstandard amino acid, p-azidophenylalanine (pAzF), at the C-terminus.^33,34^ This was subsequently conjugated to an alkyne-labeled DNA oligonucleotide with complementary sequence to the handles (termed antihandles) by copper-mediated azide/alkyne click chemistry. The conjugation product was purified by anion-exchange and size-exclusion chromatography in two consecutive steps (**Figure 1b** & **S5**–**S8**, see online methods). This conjugation method has advantages compared to conventional crosslinking chemistry such as thiol-maleimide, amine-NHS ester, or SNAP-benzylguanine, in that it cleanly allows 1:1 DNA-protein conjugation with a site-specific, single amino-acid addition that keeps the protein structure and biochemistry perturbation to a minimum. Further, it may be applied to any protein expressed by such GROs using readily available chemicals in aqueous solutions, allowing the creation of imaging standards with any fluorescent protein. The purified mEGFP-antihandle conjugate was then hybridized to the 6hb structures bearing 5–100 handles to generate desired mEGFP standards, which were purified by polyethylene glycol fractionation.^35^ These mEGFP-labeled structures were first characterized by a quantitative electrophoresis analysis. Each structure’s band mobility corresponded well to the designed numbers of mEGFP per structure, with band intensities increasing proportionally to the number of fluorophores (**Figure 1c** & **S9**), showing no evidence of mEGFP self-quenching (**Figure S10**). Negative-stain TEM imaging produced striking micrographs of these decorated structures (**Figure 1a** & **S11**) that further confirmed the expected mEGFP density and location on the 6hb tubes.

To demonstrate the utility of our origami-based brightness standards, we used them to quantify dnaC in *B. subtilis*. DnaC is a well-studied helicase that has been shown to assemble into a homo-hexameric ring at the replication fork.^36–39^ Because bacteria contain a single chromosome with two replication forks, dnaC-mEGFP puncta should appear to have 6 or 12 monomeric dnaC, depending on the proximity of the replication forks.^39^ Cells expressing dnaC-mEGFP and each of the origami standards were immobilized on separate agar pads and imaged with a widefield fluorescence microscope under the same conditions. After subtracting background fluorescence from agar pads and cell autofluorescence, spots were then picked using the ImageJ plugin MicrobeJ^40^ (**Figure 1c**–**e** & **S12**–**14**). To reduce imaging artifacts, we selected only DNA-origami spots that coincided with slightly elongated Alexa Fluor 647 spots, and dnaC spots that resided within the rod-shaped cells. Spots from origami structures were used to create a standard curve correlating fluorescence intensity to molecule number, which showed excellent linearity (**Figure 1e**), similar to the bulk measurement by gel electrophoresis (**Figure S9**). The variance of brightness from the standard structure is consistent with the heterogeneity of fluorescent output of mEGFP molecules^41^ and mEGFP labeling efficiency. In order to normalize for sub-stoichiometric labeling of DNA-origami structures, stepwise photobleaching was performed on a 6hb tube designed to carry 5 molecules of Alexa Fluor 488. Fitting the step counts to a binomial distribution yielded the fluorophore attachment probability of ∼0.80 (**Figure S15**); e.g., 100× mEGFP standard had an average of 80 fluorescent proteins, 70× standard had 56, and so on. The labeling efficiency we measured is consistent with previous reports.^23,42^ Finally, the distribution of intensities from bacterial mEGFP puncta were fit to a sum of two Gaussians and calibrated against the standard curve to derive the dnaC stoichiometry. Our method resulted in 5.50±1.97 and 11.6±2.94 (mean±SD) dnaC per puncta (**Figure 1e**), which agrees well with the expected dnaC counts of 6 and 12 molecules.^39^

In addition to quantifying widefield images of GFP-tagged protein in bacteria, we sought to demonstrate our system’s broad applications by counting dye-tagged clathrin light chain (CLC) molecules in mammalian cells using confocal microscopy (**Figure 2**). During receptor-mediated endocytosis, CLC molecules assemble with adaptors and other regulatory proteins into clathrin-coated pits and plaques, distinguished by size, clathrin number, and dynamics.^43,44^ Coated pits are smaller and more circular than plaques, making it relatively straightforward to distinguish between the two in micrographs. Coated pits assemble into clathrin cages containing varying numbers of triskelia, each comprised of 3 clathrin heavy chains and 3 clathrin light chains.^45^ The reported numbers of CLCs in a single vesicle vary widely between different tissues and methods of estimation.^44,46,47^ The recruitment, assembly and disassembly of membrane-coating clathrin structures are highly dynamic (∼45–200-s life-time; 60–100 triskelia, or ∼180–300 CLCs, for a mature clathrin-coated endocytic pit or vesicle).^44^

**Figure 2.**
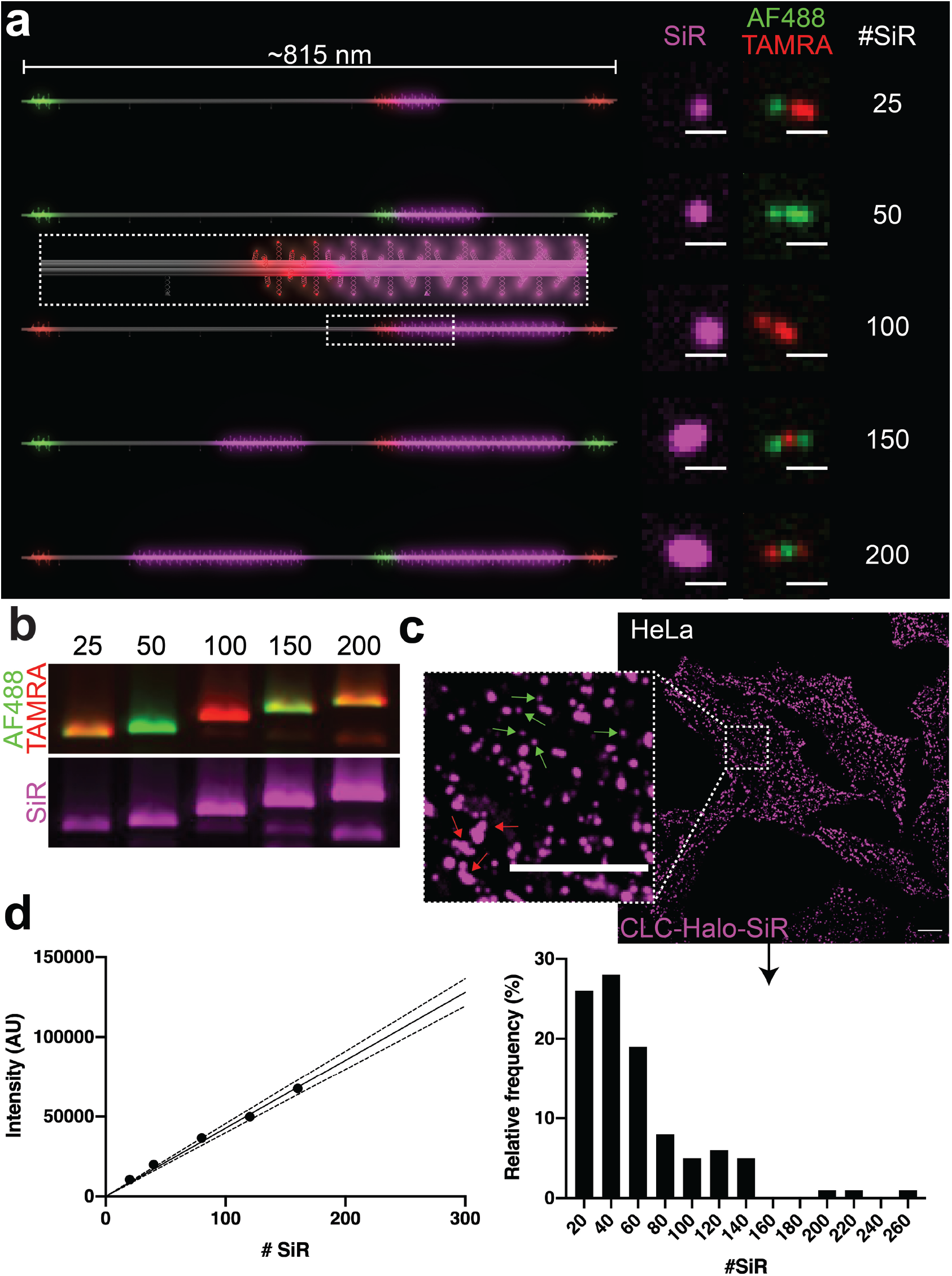
A DNA-origami-based SiR brightness standard. (a) 3D models of dimeric DNA-origami nanotubes hosting 25–200 SiR molecules (magenta) in the main body, as well as Alexa Fluor 488 (Green) and TAMRA (red) at three distinct locations for barcoding. Confocal microscopy images revealed the expected barcoding patterns and corresponding increase in SiR intensity. Scale bars: 1 µm. (b) Agarose gel images of SiR standards show the expected combinations of barcoding dyes Alexa Fluor 488 and TAMRA, as well as increasing SiR intensity. (c) Confocal images of HeLa cells expressing Halo-fused clathrin light chain (CLC) after labeling with SiR-chloroalkane. Inset shows details of coated pits (small and round, green arrows) and plaques (larger and irregularly shaped, red arrows). Scale bars: 10 µm. (d) Quantifying SiR-labeled CLCs. Left: Calibration curve generated from DNA-origami-based SiR standards (SEM too small to see). Dotted lines denote 95% confidence interval. Right: Spots containing SiR-labeled CLCs binned by molecule number.

To accommodate more fluorophores, we built dimeric 6hb nanotube structures that can in theory host up to ∼300 fluorophores (though a maximum of 200 were used in this study). The dimeric nanotube is designed to have 3 barcoding zones reserved for Alexa Fluor 488 or TAMRA (12 fluorophores per zone) to enable the selection of intact dimer structures and, potentially, sorting after simultaneous acquisition^22^ (**Figure 2a**). To quantify Halo-tagged CLC labeled with far-red dye silicon rhodamine (SiR)^48,49^ in live HeLa cells, we labeled our fluorescent standards with the same dye. Similar to the mEGFP-DNA conjugation, SiR-azide was conjugated to alkyne-DNA *via* copper-mediated click chemistry^34,50,51^ and then purified using denaturing polyacrylamide gel electrophoresis (PAGE) (**Figure S16**, also see online methods). Upon hybridizing SiR-labeled anti-handles to the DNA nanotubes, we analyzed the SiR-labeled standards using agarose gel electrophoresis, which showed SiR intensity increasing linearly with the designed dye numbers and the expected barcoding dye combinations (**Figure 2b**). The biotinylated SiR standards were then purified by rate-zonal centrifugation, individually immobilized on streptavidin-coated glass-bottom dishes, and imaged under confocal microscope (**Figure 2a** & **S17**–**22**), which confirmed high-quality dimeric structures by fluorescent barcode patterns. Next, we applied these standards to counting SiR-labeled CLCs near the surface-adhering membranes of live HeLa cells (**Figure 2c**–**d, S23**). Cells and standards were imaged on the same day under identical conditions, and SiR-fluorescent spots were selected from the background-subtracted images using a custom TrackMate^52^ script (**Figure 2d**, left & **S24**). SiR puncta in HeLa cells were manually picked to identify those resembling clathrin-coated pits (round-shaped puncta < ∼1 µm^2^), and CLC counts per cluster were quantified using our DNA-origami calibration curve (**Figure 2d**, right). Out of the 100 clusters, 97 of them contain no more than 150 CLC (median = 47.8), with a decreasing frequency as molecule number increased. This is to be expected from clathrin cages that pinch off shortly after completing assembly.^44,46^ The CLC counts obtained here is likely an underestimate because of the less-than-perfect SiR-labeling efficiency in cells. The few cellular puncta above 150 molecules are presumed to be either accidental selection of clathrin patches, or more than one pit close together.

In summary, our DNA-origami based brightness standards provide a versatile, easy-to-implement system for quantifying 5–300 clustered proteins using fluorescence microscopes readily accessible to cell biology labs. The most valuable features of our standards are their programmability to accommodate fluorophores of various types and stoichiometry, their robustness to withstand near-physiological conditions,^53^ and their low entry barrier to use without specialized equipment or software. Typically, imaging these standards adds less than 2 hours to a typical cell imaging session (compared to several days using alternative methods), and thus can be easily integrated into biologists’ conventional workflows to obtain very good estimates of protein counts. While we demonstrate the standards’ efficacy in bacteria and mammalian cells, they should, in principle, be compatible with any cell type. Similarly, because of the large library of DNA-fluorophore conjugation chemistry,^54^ including our GRO-enabled protein conjugation, our system can be used to display and count practically any fluorophore. The primary limitation of our system is that the standards are external to the cell, and so must be imaged in media similar to the cellular compartment being studied. Because the main media effectors of fluorescence output are solvent polarity and pH,^55–58^ this obstacle can be overcome by matching the pH of the aqueous imaging media to the cellular pH, as is the case in *HeLa* and *B. subtilis* cytosol.^59,60^ In cases where the cellular pH is unknown or dynamic, our universal conjugation method allows attachment of pH-sensitive fluorophores like pHluorin^55,58^ to account for any pH differences. In this work, we used a few existing, commonly used image-analysis software to extract quantitative data from micrographs. However, we envision that a specialized particle-tracking and pattern-recognition software, perhaps with future development of machine learning, will further expedite the workflow.

## Acknowledgements

We thank M. Amiram and B. Akhuetie-Oni for helpful discussions of the GRO in our conjugation system. This study is supported by a National Institutes of Health (NIH) Director’s Innovator Award (DP2-GM114830), an NIH grant (R01-GM132114), and a Yale University faculty startup fund to C.L., an NIH grant to E.K. (R01-NS113236), NIH grants (R01-GM115636 and R21-GM132661) to J.B., an NIH grant (R01-GM125951) to F.J.I., an NIH training grant (T32-EB09941) to N.D.W., and an National Science Foundation Graduate Fellowship to J.T.P.

## References

(1) Bacia, K.; Schwille, P. A Dynamic View of Cellular Processes by in Vivo Fluorescence Auto- and Cross-Correlation Spectroscopy. Methods (San Diego, Calif.) 2003, 29 (1), 74–85. https://doi.org/10.1016/s1046-2023(02)00291-8.

(2) Das, S. K.; Darshi, M.; Cheley, S.; Wallace, M. I.; Bayley, H. Membrane Protein Stoichiometry Determined from the Step-Wise Photobleaching of Dye-Labelled Subunits. 2007, 8 (9), 994–999. https://doi.org/10.1002/cbic.200600474.

(3) Ulbrich, M. H.; Isacoff, E. Y. Subunit Counting in Membrane-Bound Proteins. Nature methods 2007, 4 (4), 319–321. https://doi.org/10.1038/nmeth1024.

(4) Bulseco, D. A.; Wolf, D. E. Fluorescence Correlation Spectroscopy: Molecular Complexing in Solution and in Living Cells, 4th ed.; Digital Microscopy; Elsevier Inc., 2013; Vol. 114. https://doi.org/10.1016/b978-0-12-407761-4.00021-x.

(5) Verdaasdonk, J. S.; Lawrimore, J.; Bloom, K. Determining Absolute Protein Numbers by Quantitative Fluorescence Microscopy; Quantitative Imaging in Cell Biology; Elsevier, 2014; Vol. 123, pp 347–365. https://doi.org/10.1016/b978-0-12-420138-5.00019-7.

(6) Deatherage, C. L.; Nikolaus, J.; Karatekin, E.; Burd, C. G. Retromer Forms Low Order Oligomers on Supported Lipid Bilayers. J Biol Chem 2020, 295 (34), 12305–12316. https://doi.org/10.1074/jbc.ra120.013672.

(7) Wu, J.-Q.; Pollard, T. D. Counting Cytokinesis Proteins Globally and Locally in Fission Yeast. Science (New York, N.Y.) 2005, 310 (5746), 310–314. https://doi.org/10.1126/science.1113230.

(8) Berro, J.; Pollard, T. D. Synergies between Aip1p and Capping Protein Subunits (Acp1p and Acp2p) in Clathrin-Mediated Endocytosis and Cell Polarization in Fission Yeast. Molecular biology of the cell 2014, 25 (22), 3515–3527. https://doi.org/10.1091/mbc.e13-01-0005.

(9) Coffman, V.; Lee, I.-J.; Wu, J.-Q. Counting Molecules Within Cells; Biota Publishing; Biota Publishing, 2014.

(10) Akamatsu, M.; Lin, Y.; Bewersdorf, J.; Pollard, T. D. Analysis of Interphase Node Proteins in Fission Yeast by Quantitative and Superresolution Fluorescence Microscopy. Molecular biology of the cell 2017, 28 (23), 3203–3214. https://doi.org/10.1091/mbc.e16-07-0522.

(11) Jungmann, R.; Avendaño, M. S.; Dai, M.; Woehrstein, J. B.; Agasti, S. S.; Feiger, Z.; Rodal, A.; Yin, P. Quantitative Super-Resolution Imaging with QPAINT. Nature methods 2016, 13 (5), 439–442. https://doi.org/10.1038/nmeth.3804.

(12) Zanacchi, F. C.; Manzo, C.; Alvarez, A. S.; Derr, N. D.; Garcia-Parajo, M. F.; Lakadamyali, M. A DNA Origami Platform for Quantifying Protein Copy Number in Super-Resolution. Nature methods 2017, 14 (8), 789–792. https://doi.org/10.1038/nmeth.4342.

(13) Wassie, A. T.; Zhao, Y.; Boyden, E. S. Expansion Microscopy: Principles and Uses in Biological Research. Nature methods 2019, 16 (1), 33–41. https://doi.org/10.1038/s41592-018-0219-4.

(14) Stein, J.; Stehr, F.; Schueler, P.; Blumhardt, P.; Schueder, F.; Mücksch, J.; Jungmann, R.; Schwille, P. Towards Absolute Molecular Numbers in DNA-PAINT. Nano Letters 2019, acs.nanolett.9b03546-30. https://doi.org/10.1021/acs.nanolett.9b03546.

(15) Rothemund, P. W. K. Folding DNA to Create Nanoscale Shapes and Patterns. Nature 2006, 440 (7082), 297–302. https://doi.org/10.1038/nature04586.

(16) Wagenbauer, K. F.; Engelhardt, F. A. S.; Stahl, E.; Hechtl, V. K.; Stömmer, P.; Seebacher, F.; Meregalli, L.; Ketterer, P.; Gerling, T.; Dietz, H. How We Make DNA Origami. ChemBioChem 2017, 18 (19), 1873–1885. https://doi.org/10.1002/cbic.201700377.

(17) Bila, H.; Kurisinkal, E. E.; Bastings, M. M. C. Engineering a Stable Future for DNA-Origami as a Biomaterial. Biomaterials science 2019, 7 (2), 532–541. https://doi.org/10.1039/c8bm01249k.

(18) Fan, S.; Wang, D.; Kenaan, A.; Cheng, J.; Cui, D.; Song, J. Create Nanoscale Patterns with DNA Origami. Small 2019, 15 (26), e1805554. https://doi.org/10.1002/smll.201805554.

(19) Schmied, J. J.; Raab, M.; Forthmann, C.; Pibiri, E.; Wünsch, B.; Dammeyer, T.; Tinnefeld, P. DNA Origami–Based Standards for Quantitative Fluorescence Microscopy. Nature protocols 2014, 9 (6), 1367–1391. https://doi.org/10.1038/nprot.2014.079.

(20) Mathieu, F.; Liao, S.; Kopatsch, J.; Wang, T.; Mao, C.; Seeman, N. C. Six-Helix Bundles Designed from DNA. Nano Lett 2005, 5 (4), 661–665. https://doi.org/10.1021/nl050084f.

(21) Douglas, S. M.; Chou, J. J.; Shih, W. M. DNA-Nanotube-Induced Alignment of Membrane Proteins for NMR Structure Determination. Proceedings of the National Academy of Sciences of the United States of America 2007, 104 (16), 6644–6648. https://doi.org/10.1073/pnas.0700930104.

(22) Lin, C.; Jungmann, R.; Leifer, A. M.; Li, C.; Levner, D.; Church, G. M.; Shih, W. M.; Yin, P. Submicrometre Geometrically Encoded Fluorescent Barcodes Self-Assembled from DNA. Nature Chemistry 2012, 4 (10), 832–839. https://doi.org/10.1038/nchem.1451.

(23) Schmied, J. J.; Gietl, A.; Holzmeister, P.; Forthmann, C.; Steinhauer, C.; Dammeyer, T.; Tinnefeld, P. Fluorescence and Super-Resolution Standards Based on DNA Origami. Nature methods 2012, 9 (12), 1133–1134. https://doi.org/10.1038/nmeth.2254.

(24) Schmied, J. J.; Forthmann, C.; Pibiri, E.; Lalkens, B.; Nickels, P.; Liedl, T.; Tinnefeld, P. DNA Origami Nanopillars as Standards for Three-Dimensional Superresolution Microscopy. Nano Letters 2013, 13 (2), 781–785. https://doi.org/10.1021/nl304492y.

(25) Jusuk, I.; Vietz, C.; Raab, M.; Dammeyer, T.; Tinnefeld, P. Super-Resolution Imaging Conditions for Enhanced Yellow Fluorescent Protein (EYFP) Demonstrated on DNA Origami Nanorulers. Scientific Reports 2015, 1–9. https://doi.org/10.1038/srep14075.

(26) Woehrstein, J. B.; Strauss, M. T.; Ong, L. L.; Wei, B.; Zhang, D. Y.; Jungmann, R.; Yin, P. Sub-100-Nm Metafluorophores with Digitally Tunable Optical Properties Self-Assembled from DNA. Science Advances 2017, 3 (6). https://doi.org/10.1126/sciadv.1602128.

(27) Werner, E. W.; Mei, T.-S.; Burckle, A. J.; Sigman, M. S. Enantioselective Heck Arylations of Acyclic Alkenyl Alcohols Using a Redox-Relay Strategy. Science (New York, N.Y.) 2012, 338 (6113), 1455–1458. https://doi.org/10.1126/science.1229208.

(28) Czogalla, A.; Petrov, E. P.; Kauert, D. J.; Uzunova, V.; Zhang, Y.; Seidel, R.; Schwille, P. Switchable Domain Partitioning and Diffusion of DNA Origami Rods on Membranes. Faraday Discuss. 2013, 161, 31–43. https://doi.org/10.1039/c2fd20109g.

(29) Douglas, S. M.; Dietz, H.; Liedl, T.; Högberg, B.; Graf, F.; Shih, W. M. Self-Assembly of DNA into Nanoscale Three-Dimensional Shapes. Nature 2009, 459 (7245), 414–418. https://doi.org/10.1038/nature08016.

(30) Lin, C.; Perrault, S. D.; Kwak, M.; Graf, F.; Shih, W. M. Purification of DNA-Origami Nanostructures by Rate-Zonal Centrifugation. Nucleic acids research 2013, 41 (2), e40–e40. https://doi.org/10.1093/nar/gks1070.

(31) Bellot, G.; McClintock, M. A.; Chou, J. J.; Shih, W. M. DNA Nanotubes for NMR Structure Determination of Membrane Proteins. Nature protocols 2013, 8 (4), 755–770. https://doi.org/10.1038/nprot.2013.037.

(32) Stahl, E.; Martin, T. G.; Praetorius, F.; Dietz, H. Facile and Scalable Preparation of Pure and Dense DNA Origami Solutions. Angewandte Chemie International Edition 2014, 53 (47), 12735–12740. https://doi.org/10.1002/anie.201405991.

(33) Lajoie, M. J.; Rovner, A. J.; Goodman, D. B.; Aerni, H.-R.; Haimovich, A. D.; Kuznetsov, G.; Mercer, J. A.; Wang, H. H.; Carr, P. A.; Mosberg, J. A.; Rohland, N.; Schultz, P. G.; Jacobson, J. M.; Rinehart, J.; Church, G. M.; Isaacs, F. J. Genomically Recoded Organisms Expand Biological Functions. Science (New York, N.Y.) 2013, 342 (6156), 357–360. https://doi.org/10.1126/science.1241459.

(34) Amiram, M.; Haimovich, A. D.; Fan, C.; Wang, Y.-S.; Aerni, H.-R.; Ntai, I.; Moonan, D. W.; Ma, N. J.; Rovner, A. J.; Hong, S. H.; Kelleher, N. L.; Goodman, A. L.; Jewett, M. C.; Söll, D.; Rinehart, J.; Isaacs, F. J. Evolution of Translation Machinery in Recoded Bacteria Enables Multi-Site Incorporation of Nonstandard Amino Acids. Nature biotechnology 2015, 33 (12), 1272–1279. https://doi.org/10.1038/nbt.3372.

(35) Shaw, A.; Benson, E.; Högberg, B. Purification of Functionalized DNA Origami Nanostructures. Acs Nano 2015, 9 (5), 4968–4975. https://doi.org/10.1021/nn507035g.

(36) Fass, D.; Bogden, C. E.; Berger, J. M. Crystal Structure of the N-Terminal Domain of the DnaB Hexameric Helicase. Structure 1999, 7 (6), 691–698. https://doi.org/10.1016/s0969-2126(99)80090-2.

(37) Bailey, S.; Eliason, W. K.; Steitz, T. A. The Crystal Structure of the Thermus Aquaticus DnaB Helicase Monomer. Nucleic acids research 2007, 35 (14), 4728–4736. https://doi.org/10.1093/nar/gkm507.

(38) Kaplan, D. L.; Saleh, O. A.; Ribeck, N. Single-Molecule and Bulk Approaches to the DnaB Replication Fork Helicase. Frontiers in Bioscience-Landmark 2013, 18 (1), 224–240. https://doi.org/10.2741/4097.

(39) Mangiameli, S. M.; Merrikh, C. N.; Wiggins, P. A.; Elife, H. M.; 2017. Transcription Leads to Pervasive Replisome Instability in Bacteria. elifesciences.org n.d. https://doi.org/10.7554/elife.19848.001.

(40) Ducret, A.; Quardokus, E. M.; Brun, Y. V. MicrobeJ, a Tool for High Throughput Bacterial Cell Detection and Quantitative Analysis. 2016, 1–7. https://doi.org/10.1038/nmicrobiol.2016.77.

(41) Tsien, R. Y. The Green Fluorescent Protein. Annual review of biochemistry 1998, 67 (1), 509–544. https://doi.org/10.1146/annurev.biochem.67.1.509.

(42) Acuna, G. P.; Möller, F. M.; Holzmeister, P.; Beater, S.; Lalkens, B.; Tinnefeld, P. Fluorescence Enhancement at Docking Sites of DNA-Directed Self-Assembled Nanoantennas. Science (New York, N.Y.) 2012, 338 (6), 506. https://doi.org/10.1126/science.1228638.

(43) Saffarian, S.; Cocucci, E.; Kirchhausen, T. Distinct Dynamics of Endocytic Clathrin-Coated Pits and Coated Plaques. PLOS Biology 2009, 7 (9), e1000191–18. https://doi.org/10.1371/journal.pbio.1000191.

(44) Sochacki, K. A.; Taraska, J. W. From Flat to Curved Clathrin: Controlling a Plastic Ratchet. Trends in Cell Biology 2019, 29 (3), 241–256. https://doi.org/10.1016/j.tcb.2018.12.002.

(45) Harris, J. R.; Marles-Wright, J. Macromolecular Protein Complexes; Springer; Springer, 2017.

(46) Ehrlich, M.; Boll, W.; Oijen, A. van; Hariharan, R.; Chandran, K.; Nibert, M. L.; Kirchhausen, T. Endocytosis by Random Initiation and Stabilization of Clathrin-Coated Pits. Cell 2004, 118 (5), 591–605. https://doi.org/10.1016/j.cell.2004.08.017.

(47) Otter, W. K. den; Renes, M. R.; Briels, W. J. Asymmetry as the Key to Clathrin Cage Assembly. Biophysical journal 2010, 99 (4), 1231–1238. https://doi.org/10.1016/j.bpj.2010.06.011.

(48) Lukinavicius, G.; Umezawa, K.; Olivier, N.; Honigmann, A.; Yang, G.; Plass, T.; Mueller, V.; Reymond, L.; Jr, I. R. C.; Luo, Z.-G.; Schultz, C.; Lemke, E. A.; Heppenstall, P.; Eggeling, C.; Manley, S.; Johnsson, K. A Near-Infrared Fluorophore for Live-Cell Super-Resolution Microscopy of Cellular Proteins. Nature Chemistry 2013, 5 (2), 132–139. https://doi.org/10.1038/nchem.1546.

(49) Thompson, A. D.; Omar, M. H.; Rivera-Molina, F.; Xi, Z.; Koleske, A. J.; Toomre, D. K.; Schepartz, A. Long-Term Live-Cell STED Nanoscopy of Primary and Cultured Cells with the Plasma Membrane HIDE Probe DiI-SiR. Angewandte Chemie International Edition 2017, 56 (35), 10408–10412. https://doi.org/10.1002/anie.201704783.

(50) El-Sagheer, A. H.; Brown, T. Click Chemistry with DNA. Chemical Society Reviews 2010, 39 (4), 1388–19. https://doi.org/10.1039/b901971p.

(51) Presolski, S. I.; Hong, V. P.; Finn, M. G. Copper-Catalyzed Azide-Alkyne Click Chemistry for Bioconjugation. Current protocols in chemical biology 2011, 3 (4), 153–162. https://doi.org/10.1002/9780470559277.ch110148.

(52) Tinevez, J.-Y.; Perry, N.; Schindelin, J.; Hoopes, G. M.; Reynolds, G. D.; Laplantine, E.; Bednarek, S. Y.; Shorte, S. L.; Eliceiri, K. W. TrackMate: An Open and Extensible Platform for Single-Particle Tracking. Methods (San Diego, Calif.) 2017, 115, 80–90. https://doi.org/10.1016/j.ymeth.2016.09.016.

(53) Kielar, C.; Xin, Y.; Shen, B.; Kostiainen, M. A.; Grundmeier, G.; Linko, V.; Keller, A. On the Stability of DNA Origami Nanostructures in Low-Magnesium Buffers. Angewandte Chemie International Edition 2018, 57 (30), 9470–9474. https://doi.org/10.1002/anie.201802890.

(54) Yang, Y. R.; Liu, Y.; Yan, H. DNA Nanostructures as Programmable Biomolecular Scaffolds. Bioconjugate chemistry 2015, 26 (8), 1381–1395. https://doi.org/10.1021/acs.bioconjchem.5b00194.

(55) Miesenböck, G.; Angelis, D. A. D.; Rothman, J. E. Visualizing Secretion and Synaptic Transmission with PH-Sensitive Green Fluorescent Proteins. Nature 1998, 394 (6689), 192–195. https://doi.org/10.1038/28190.

(56) Straight, A. F. Fluorescent Protein Applications in Microscopy; Digital Microscopy, 3rd Edition; Elsevier, 2007; Vol. 81, pp 93–113. https://doi.org/10.1016/s0091-679x(06)81006-x.

(57) Papkovsky, D. Live Cell Imaging; Humana Press; Humana Press, 2009.

(58) Mahon, M. J. PHluorin2: An Enhanced, Ratiometric, PH-Sensitive Green Florescent Protein. Advances in bioscience and biotechnology (Print) 2011, 2 (3), 132–137. https://doi.org/10.4236/abb.2011.23021.

(59) Llopis, J.; McCaffery, J. M.; Miyawaki, A.; Farquhar, M. G.; Tsien, R. Y. Measurement of Cytosolic, Mitochondrial, and Golgi PH in Single Living Cells with Green Fluorescent Proteins. Proceedings of the National Academy of Sciences of the United States of America 1998, 95 (12), 6803–6808. https://doi.org/10.1073/pnas.95.12.6803.

(60) Beilen, J. W. A. van; Brul, S. Compartment-Specific PH Monitoring in Bacillus Subtilis Using Fluorescent Sensor Proteins: A Tool to Analyze the Antibacterial Effect of Weak Organic Acids. Frontiers in Microbiology 2013, 4. https://doi.org/10.3389/fmicb.2013.00157.

