## Supplementary Information for "DNA-Origami-Based Fluorescence Brightness Standards for Convenient and Fast Protein Counting in Live Cells"

### Materials and Methods

#### Materials

DNA oligonucleotides were synthesized by Integrated DNA Technologies. All chemical reagents were purchased from Millipore Sigma unless otherwise specified. Enzymes were purchased from New England Biolabs unless otherwise specified. Water is purified by a Milli-Q (Millipore) system.

#### Agarose gel electrophoresis

Unless otherwise indicated, all agarose gels contained 1.5% agarose, and were run in 0.5×TBE + 10 mM $MgCl_2$ for 3 hrs at 5V/cm at room temperature. Gels were imaged on a Typhoon FLA 9500 scanner. Whenever necessary, gels were stained with Ethidium Bromide (EtBr, 0.5 µg/mL) after imaging any other relevant channels (for fluorescent labels on DNA-origami structures).

#### SDS-polyacrylamide gel electrophoresis (SDS-PAGE)

Unless otherwise indicated, all gels contained 12% acrylamide bis-tris (Bio-Rad), and were run for 45 minutes at 200 V in 1× SDS MOPS buffer (50 mM Tris base, 50 mM MOPS, 1 mM EDTA, 0.1% SDS, pH 6.5). Samples were prepared in 30% glycerol supplemented with orange G dye, and were heated 5 minutes at 95ºC before loading. In gels where mEGFP was imaged, SDS was first rinsed away by three 20-minute washes in water. We found this protocol was able to recover mEGFP fluorescence sufficiently to image (See Figure S5).
SYBR Gold staining was performed by submerging gels in SYBR Gold stain (Invitrogen) diluted 10,000× in water, per manufacturer recommendations. Gels were briefly rinsed with water before imaging on a Typhoon FLA 9500 scanner.
Coomassie staining was performed by submerging gels in 1× Coomassie solution (50% methanol, 10% glacial acetic acid, 0.1% w/v Coomassie blue) and microwaving on high until boiling (approximately 1 minute). Gels were then briefly rinsed with water, covered in Coomassie destain solution (12% methanol, 7% glacial acetic acid), and microwaved on high until boiling (approximately 1 minute). Containers with gels heated in destain solution were then placed on a platform shaker, and Kimwipes were added to speed Coomassie desorption. Gels were checked approximately every 30 minutes until clear enough to be imaged. Coomassie-stained gels were imaged on a transilluminatior (Bio-Rad) with white light. In cases where multiple scans were used, gels were imaged and stained in the following order: in-gel fluorophores (e.g. mEGFP), SYBR Gold, and Coomassie.

#### DNA-origami structure design

DNA-origami six-helix bundle (6hb) nanotubes were designed using caDNAno (cadnano.org)[1]. Design diagrams are shown in Figure S4. Orthogonal handle sequences were generated using NUPACK (nupack.org) and added to the 3'-ends of selected staple strands. Handle sequences are shown in Table S1.

#### DNA-origami structure folding and purification

DNA origami folding mixtures were prepared by adding 6× molar excess of staple strands to p7308 scaffold[2] in 1×TE buffer (25 mM Tris•HCl, 1 mM EDTA•$Na_2$, pH 8) supplemented with 10 mM $MgCl_2$. Mixtures were heated in a thermocycler (Bio-Rad) to 85ºC for 3 min, cooled from 80ºC to 60ºC in 80 minutes, 60ºC to 24ºC in 15 hours, and then held at 4ºC. Excess staples were removed by PEG precipitation and re-suspending pellets in 1×TE buffer + 10 mM $MgCl_2$.[3]

**DNA-origami structure dimerization**
"Front" and "rear" monomeric halves of dimeric nanotubes were folded independently and PEG precipitated to remove excess staples (see above). Concentrations of purified monomers were estimated by absorbance, and equimolar amounts of each half were mixed in 1×TE buffer + 10 mM $MgCl_2$ with 10× molar excess of linker strands (Table S2). These dimer mixtures were heated to 55ºC and cooled to 20ºC over 18 hours.

**Fluorescently labeling DNA-origami structures**
To add antihandles to DNA nanotubes, a molar excess of antihandle was added to origami in 1×TE buffer + 10 mM $MgCl_2$: 3× excess for mEGFP and SiR antihandles, and 1.2× excess in the case of other antihandles. 3× excess was used to maximize antihandle attachment for the most important fluorophores. The mixture is incubated at 37ºC for 2 hours. In the case of dimeric structures, linker DNA was kept at 10× molar excess during the labeling step. The labeled DNA-origami dimers were then purified by rate-zonal centrifugation (see below).

**mEGFP standard purification**
DNA-origami nanotubes labeled by mEGFP and Alexa Fluor 647 were PEG precipitated to remove excess antihandles. Briefly, a equal volume of 2×PEG-precipitation buffer (1×TE, 15% PEG, 10 mM $MgCl_2$, 500 mM NaCl, pH 8) was added to the hybridization mixture (DNA nanotubes + labeled-antihandles), and the mixture was spun at 16,000-g for 25 minutes at 4°C. The pellet was then resuspended in 1×TE + 10 mM $MgCl_2$. A typical preparation starting with 45 µL of 4 nM unlabeled DNA nanotubes yields 45 µL of 2.7 nM purified mEGFP-labeled DNA nanotubes, over 67% recovery.

**Transmission electron microscopy**
For negative-stain TEM, a drop of the sample (5 µL) was deposited on a glow discharged formvar/carbon-coated copper grid (Electron Microscopy Sciences), incubated for 1 min, and blotted away. The grid was first rinsed twice with 1× TE buffer + 10 mM $MgCl_2$, then washed briefly and stained for 1 min with 2% (w/v) uranyl formate. Images were acquired on a JEOL JEM-1400Plus microscope (acceleration voltage: 80 kV) with a bottom-mount 4k×3k CCD camera (Advanced Microscopy Technologies).

**SiR standard purification**
Dimeric DNA-origami nanotubes labeled with fluorophores and biotin were purified by rate-zonal centrifugation over a glycerol gradient[4]. Gradients were 15%–45% glycerol in 1×TE buffer supplemented with 10 mM $MgCl_2$, and were spun at 48k RPM for 1.5 hrs at 4ºC. Fractions were subsequently collected and run in agarose gels to find the fractions

containing properly labeled dimers. A typical preparation starting with 90 µL of 15 nM dimeric DNA nanotubes yields 600 µL of ~1 nM purified SiR-labeled DNA nanotubes.

**mEGFP-pAzF cloning**
The sequence for mEGFP was cloned from plasmid pFA6a-mEGFP-kanMX6 (Addgene plasmid # 87023; RRID:Addgene_87023), and was inserted into destination vector pZE21-GFP-NHis-0TAG generated by the Isaacs lab[5]. First, overlap-extension PCR was used to add an EcoRI restriction site and a 6×His tag to the N-terminal of mEGFP, and a TAG codon and BamHI restriction site to the C-terminus. The resulting PCR product and destination vector were both digested with EcoRI and BamHI and purified by agarose gel electrophoresis. The gel-purified products were then ligated using T4 ligase. The final plasmid introduced into the GRO contained mEGFP with a 6×His N-term tag and a C-term TAG codon with inducible expression controlled by a pLtetO promoter. DH5α cells were transformed with this plasmid and plated on kanamycin-selective agar plates. 20 colonies were picked and PCR tested with the forward and reverse primers used to clone the mEGFP insert. 4 colonies that produced the expected PCR result were grown overnight and extracted using a Miniprep kit (Qiagen). The resulting plasmid DNA was sequenced. Of the 4 sequenced colonies, 1 contained the exact 6×His-mEGFP-TAG sequence as designed. A small volume of the DH5α culture with the correct mEGFP was stored at -80ºC in 25% glycerol.

**GRO transformation**
Recoded E. coli ΔmutS:Zeo (Δ1prfA):tolC)tolC[5,6] was transformed with the plasmid 6×His_mEGFP-TAG (see above) and orthogonal translation system plasmid pAzFRS.1.t1[5], both generated by the Isaacs lab, and plated on agar containing chloramphenicol and kanamycin. A resulting colony was picked and used to grow a ~5 mL culture, ~1 mL of which was stored at -80ºC in 25% glycerol.

**mEGFP-pAzF overexpression** (adapted from ref 6)
A 100 mL 2×YT starter culture supplemented with chloramphenicol and kanamycin was inoculated with a miniscule volume from the glycerol stock of GRO cells transformed with 6×His_mEGFP-TAG and pAzFRS.1.t1 plasmids and grown at 34ºC overnight. This starter culture was used to inoculate a 1L expression culture supplemented with chloramphenicol, kanamycin, pAzF, and arabinose. The expression culture was grown at 34ºC to confluency ($OD_{600}$ = 0.5–0.8), and then mEGFP expression was induced with addition of anhydrotetracycline (aTc). The induced culture was grown for ~16 hours at 34ºC. Then cells were pelleted by centrifugation at 4,000× g for 15 minutes. Supernatant was discarded and cell pellets were flash frozen and stored at -80ºC.

**mEGFP(pAzF) His-trap FPLC purification**
Cell pellets were thawed on ice and resuspended in lysis/wash buffer (1× PBS pH 7.4 + 25 mM imidazole, supplemented with a protease inhibitor cocktail (Roche)). Resuspended cells were lysed using a homogenizer, lysate was centrifuged at 35,000 RPM for 45 minutes, and supernatant was collected and filtered using 0.45 µm filters. A HisTrap column was conditioned with lysis/wash buffer, and then filtered lysate was circulated ~2.5× over column to bind. The column was then washed with ~3 column

volumes of lysis/wash buffer, which was collected. Bound protein was then eluted with elution buffer (1× PBS pH 7.4 + 500 mM imidazole), and fractions were collected. Fractions most likely to contain the desired product were selected by reviewing the FPLC traces (absorption at 254 & 280 nm) and were run in a reducing 12% Bis-Tris SDS-PAGE gel. Fractions confirmed to contain the desired products were pooled and buffer exchanged into lysis/wash buffer using Amicon 10k filters, then purified a second time using the HisTrap column. Fractions from the second HisTrap purification were run on a SDS-PAGE gel as before to identify the best fractions, which were then pooled, buffer exchanged into 1×PBS, pH 7.4, and measured for concentration using a BCA assay (Pierce). Finally, glycerol was added to the purified mEGFP(pAzF), which was then flash-frozen and stored at -80ºC.

**mEGFP – alkyne DNA conjugation and purification**

Click reaction was adapted from protocols by Presolski et al.[7] mEGFP(pAzF) was reacted with an excess of alkyne DNA (Table S1) for 1 hr at 30ºC in degassed click reaction buffer (0.1 M potassium phosphate, 0.25 mM $CuSO_4$, 1.25 mM THPTA, 5 mM aminoguanidine HCl, 5 mM sodium ascorbate, pH 7). The reaction was then quenched by adding EDTA to a final concentration of 2.5 mM. Anion exchange was then performed using Pierce Strong Anion Exchange Spin Columns following the manufacturer's directions. Fractions were collected and stored O/N at 4ºC before running on a 12% SDS-PAGE gel (Figure S6). Fractions confirmed to contain mEGFP-DNA conjugate but not the unreacted mEGFP were buffer exchanged into 1×PBS using 10k Amicon filters before size exclusion purification. Size exclusion was performed using a Superdex 75 Increase 10/300 column (Cytiva) connected to an ÄKTA FPLC system (Cytiva), and fractions were collected and stored at 4ºC O/N before checking with SDS-PAGE (Figure S7&S8).

**SiR-azide – alkyne-DNA click reaction**

SiR-azide (Spirochrome) was reacted to alkyne DNA (Table S1) using the same protocol as mEGFP conjugation (above).

**Urea-PAGE Purification of SiR-DNA conjugate.**

SiR-DNA conjugate was mixed 1:1 with denaturing tracking dye buffer (90% formamide, 10 mM NaOH, 1 mM EDTA-$Na_2$, 0.1% xylene cyanole) and heated 10 minutes at 95°C. Sample was removed from heat and immediately chilled on ice before being loaded into a denaturing polyacrylamide gel (12% acrylamide, 8.3 M urea, 89 mM tris base, 89 mM boric acid, 2 mM EDTA pH 8.0). Each gel well was loaded with 20 µL of SiR-DNA conjugate at ~2 OD, and the loaded urea-PAGE gel was run in 1× TBE buffer (89 mM tris base, 89 mM boric acid, 2 mM EDTA, pH 8.0) for 2 hours at a constant current of 30 mA and at 37°C. The gel was removed from the running apparatus, wrapped in plastic, and visually inspected on a transilluminator under white light and 302 nm UV to locate the migrated SiR-DNA conjugate. The desired SiR-DNA oligonucleotide was observable as a blue band under white light and as dark shadow cast against a phosphor screen under 302 nm UV. The desired DNA band was excised with a clean razorblade. The gel fragments containing the SiR-DNA were diced into smaller pieces and placed into Freeze 'N Squeeze filters (Bio-Rad) along with 500 µL of elution buffer (500 mM

$NH_4AC$, 10 mM $Mg(AC)_2$•$(H2O)_4$, 2 mM EDTA, pH 8.0). Freeze 'N Squeeze filters containing cut gel and elution buffer were wrapped in Al foil and agitated at room temperature for 8 hours. Filters were spun at room temperature at 8,000 rpm for 6 minutes to separate eluted SiR-DNA conjugate and gel. Eluted SiR-DNA was subsequently purified of organic contaminants via extraction with butanol (using a butanol:DNA volume ratio of 2:1). The extracted aqueous layer containing the SiR-DNA was then subjected to ethanol purification: 100% ethanol was added to the SiR-DNA solution at a 2:1 (v/v) ratio, mixed thoroughly, and then chilled at -20°C for 30 minutes. The tubes were then centrifuged at 13,000 rpm for 30 minutes at 4°C to pellet the SiR-DNA. The supernatant was decanted, and the SiR-DNA pellets were washed with ice-cold 70% ethanol followed by another 13,000-rpm spin for 10 minutes. After decanting the supernatant, the SiR-DNA pellets were air-dried overnight at room temperature in dark. After drying, solid SiR-DNA pellets were either dissolved in water/buffer of choice or stored dry at -20°C. Purification of the desired SiR-DNA fragment was confirmed by subsequent urea-PAGE analysis and comparison with Ultra Low Range DNA Ladder (Invitrogen). Importantly, the entire purification protocol was performed in dim lighting conditions with foil coverings over tubes and the gel running apparatus to minimize potential photobleaching of the SiR-DNA conjugate.

**Generation of *B. subtilis* cells expressing dnaC-mEGFP**
*B. subtilis* PY79 cells expressing dnaC-mEGFP (strain NW001) was constructed using plasmid DNA and a 1-step competence method. Original plasmid (dnaC-GFPmut2) was obtained from Alan D. Grossman at MIT Department of Biology and mutated to mEGFP using QuikChange Lightning kit (Agilent) following manufacturer's instructions. Mutations were confirmed by sequencing.

**Imaging of mEGFP standards and *B. subtilis* cells**
mEGFP-labeled DNA nanotubes were immobilized on a coverslip as previously described[8] for quality control imaging (not shown), and on a 2% agarose pad containing 1× M9 salts medium (Gibco) for generating calibration curves (Figure 1c & S12). Cells were prepared as previously described by Mangiameli et al.[9] with slight modifications. Briefly, *B. Subtilis* (strain NW001) were cultured overnight in arabinose minimal medium in a shaking incubator at 30°C. Overnight cultures at an $OD_{600}$ of 0.4–0.9 were diluted back to an $OD_{600}$ of 0.2 and incubated again for about 2 hr until they reached approximately $OD_{600}$ 0.4. Cells were mounted on a 2% agarose pad containing 1× M9 salts medium (Gibco) using a gene frame (Bio-Rad). Fluorescence microscopy was performed using a Leica DMi8 Wide-field Inverted Microscope equipped with an HC PL APO 100×DIC objective, an iXon Ultra 888 EMCCD Camera (Andor Technology) and Lumencor´s Spectra-X LED Light Engine as the source of light. Excitation light transmission was set to 50% and exposure time for GFP ($\lambda_{EX}$=470/40; $\lambda_{EM}$=500–550) was 1 sec. Cells were concentrated 10× by centrifugation (3300×g for 30 sec) prior to visualization. Cells were imaged at RT. Images were acquired with Leica Application Suite X, and analysis and processing were performed using the ImageJ software.

**Image processing of wide-field microscopy images**

Background fluorescence from agar pads and cell autofluorescence was estimated by measuring line profiles spanning cells and origami, then subtracted from the image. Spots were picked using MicrobeJ[10], and were subsequently selected manually: DNA-origami spots in the mEGFP channel were selected if they were non-overlapping and colocalized with signal in the Alexa Fluor 647 channel; dnaC spots were selected if they were clear, round puncta within cell boundaries (2 out of 77 puncta were excluded because of their abnormally large size and intensity). These intensities were plotted as a histogram and fit to a sum of two Gaussians function using GraphPad Prism[11]. The workflow is summarized in Figure S14.

**CLC (Clathrin Light Chain)-HaloTag CRISPR/Cas9 gene editing**
CLC-HaloTag CRISPR cells were generated by transfecting a repair template that has Halo tag and the gRNA specific to the targeted gene. The following gRNA sequence was used for the CLC genomic DNA (Gene ID 1211): 5'- GCAGATGTAGTGTTTCCACA GGG-3' (PAM sequence is underlined). This gRNA was cloned into the SpCas9 pX330 plasmid (Addgene plasmid #42230)[12] by BbSI site. The homologous repair plasmid with HaloTag was constructed by pEGFP-C1 plasmid. The right homology arm (~1 kb) was cloned into pEGFP-C1 using EcoO109I site. The left homology arm (~1 kb) was cloned with PCR fragment of HaloTag by In-Fusion HD Cloning kit (Takara Bio USA, Inc.) using AseI and BamHI cutting sites. The target sequence (PAM site) was mutagenized in the right homologous arm.
The pX330 plasmid with gRNA of CLC and the homologous repair plasmid were transfected in HeLa CCL-2 cells using Lipofectamine 3000 (Thermo Fisher). After 48hrs post-transfection, cells were selected by G418. After selection, cells were screened by single cell cloning with serial dilution protocol in 96 well plate and immunoblot.

**CLC-Halo cell cultures**
*Halo–CLC*[13] CRISPR/Cas9 HeLa cells were cultured in Dulbecco's modified Eagle medium (DMEM) (Gibco) supplemented with 10% FBS. All cells were cultured at 37°C in 5% $CO_2$ incubator.

**CLC-Halo labeling with SiR-chloroalkane (SiR-CA) in Hela cells**
Cells were seeded on 35 mm glass bottom dishes (Mattek P35G-1.5-14-C) 24 hrs before imaging. On the day of imaging *Halo–CLC* expressed in cells were labelled with 5 µM near far-red silicon rhodamine (SiR): SiR-chlroroalkane (a gift from Promega) for 30 min at 37°C. Subsequently the cells were washed three times and placed back in 37°C incubator for 1 hr.

**Imaging SiR standards and SiR labeled CLC-Halo in HeLa cells**
SiR-labeled DNA nanotube dimers were immobilized on a Mattek dish as previously described[8], in Live Cell Imaging Solution (Thermo Fisher, pH 7.4). The cells were imaged live on TiE inverted Nikon spinning disc confocal microscope, using a 100× 1.45 Oil objective. SiR labels were imaged using a 647 nm laser line (190 mW, measured close to the coverslip) using Nikon's Perfect Focus System. Laser power was maintained constant throughout the imaging session after adjusting (50–60%) to avoid

saturation. Several images of glass bottom dishes without cells were also captured at the same focal point for background correction post acquisition (see below).

**Image processing of confocal micrographs**

First, background from media and the cell dish was approximated by averaging 10 images of empty dishes to generate a new image. This image was then subtracted from all images containing DNA-origami nanotubes or HeLa cells. SiR puncta in DNA-origami images were picked using a custom TrackMate script (Do_subpixel_localization: True, Radius: 0.8, Threshold: 6.0, and Do_median_filtering: False), and used to construct the calibration curve (Figure 2d). A background-subtracted image of cells was median filtered (20 px radius) to generate the approximate background intensity from cell autofluorescence and nonspecific SiR-CA labeling. This image was then subtracted from the image of cells. CLC-SiR spots in cells were then picked manually by size and morphology. Only spots that were well-defined, nonoverlapping, in-focus, and circular, were selected. These criteria were selected based on the morphological differences between coated pits and plaques previously described.[14] CLC-SiR intensities were then converted to molecule number using the calibration curve. The workflow is summarized in Figure S24.

### Supplementary Figures

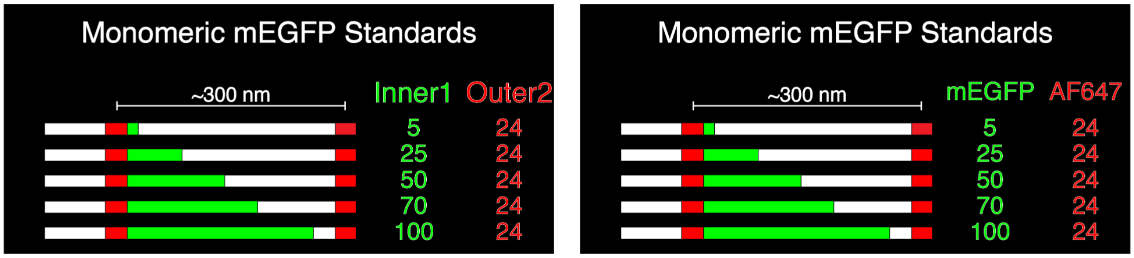

**Figure S1. Diagrams of mEGFP standards.** Left: number and names of handle sequences. Right: number and names of fluorophores.

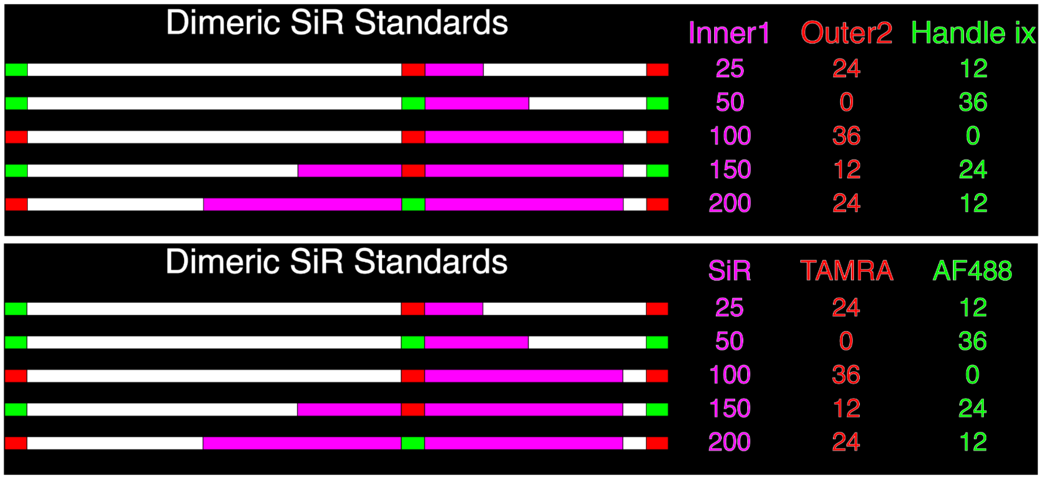

**Figure S2. Diagrams of SiR standards.** Top: number and names of handle sequences. Bottom: number and names of fluorophores.

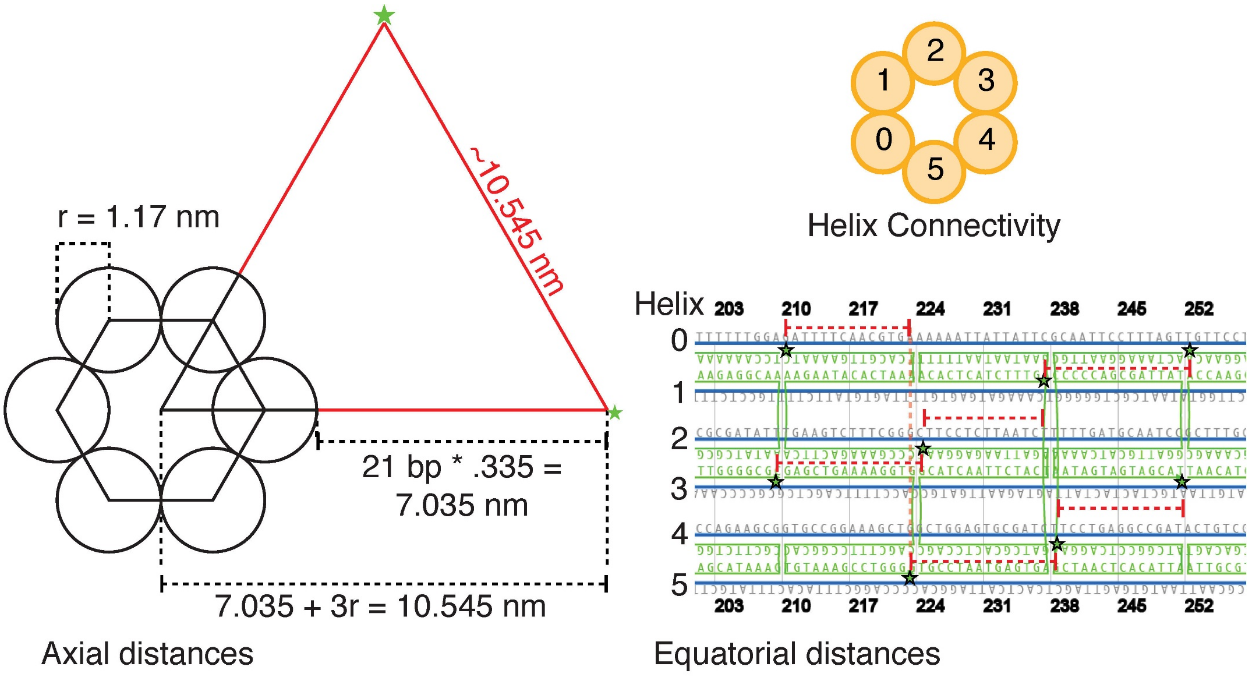

| Helix A | Helix B | Axial D (nm) | Equatorial D (NT) | D/NT (nm) | Equatorial D (nm) | Total D (nm) |
|---|---|---|---|---|---|---|
| 0 | 1 | 10.545 | 16 | 0.335 | 5.36 | 11.83 |
| 1 | 2 | 10.545 | 14 | 0.335 | 4.69 | 11.54 |
| 2 | 3 | 10.545 | 16 | 0.335 | 5.36 | 11.83 |
| 3 | 4 | 10.545 | 14 | 0.335 | 4.69 | 11.54 |
| 4 | 5 | 10.545 | 16 | 0.335 | 5.36 | 11.83 |
| 5 | 0 | 10.545 | 14 | 0.335 | 4.69 | 11.54 |

**Figure S3. Distances between fluorophores.** 21-bp handles extend from the 3' end of green staple strands (green stars). Distances between adjacent fluorophores were calculated as follows: sqrt((axial d)$^2$ + (equatorial d)$^2$). The radius of a double helix in a 6hb nanotube was measured previously.[15]

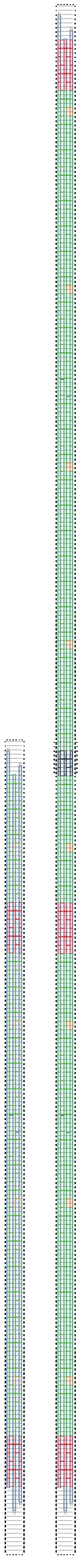

**Figure S4. Design (caDNAno[1]) diagrams of DNA-origami nanotubes**
Staples in monomeric (left) and dimeric (right) structures are color-coded. All handles extend from 3’ end.
Gray: poly-T end caps.
Red: Handles reserved for barcoding fluorophores. Handles are outer 2 or handle ix, depending on structure.
Green: optional attachment of inner 1 handle (for mEGFP/SiR).
Orange: non ATG 1 handle for biotin.
Black: linker DNA for dimerization.

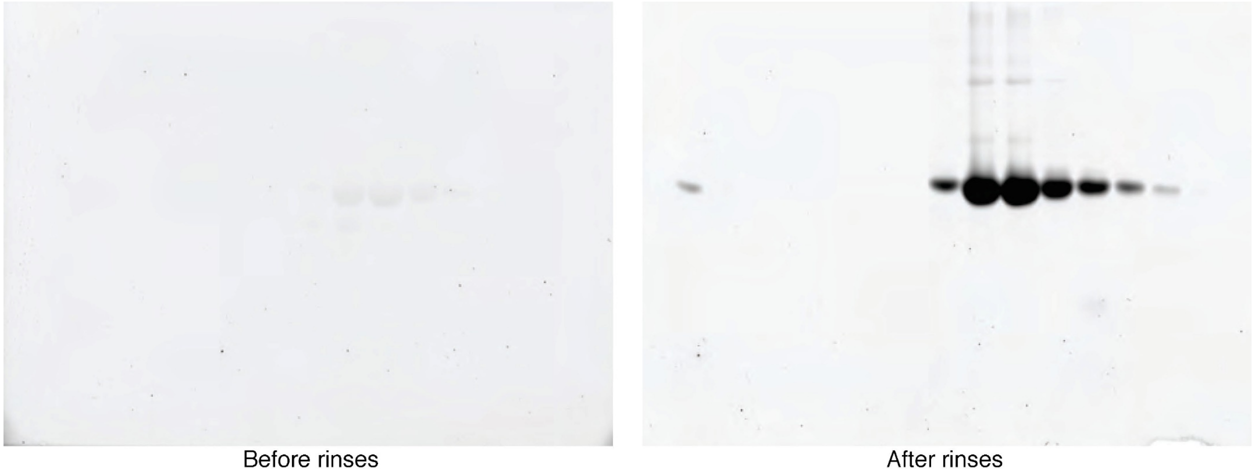

**Figure S5. mEGFP in-gel fluorescence recovery.** After rinsing in water 3× 20 min, enough fluorescence was recovered to allow gel imaging in the 488 channel.

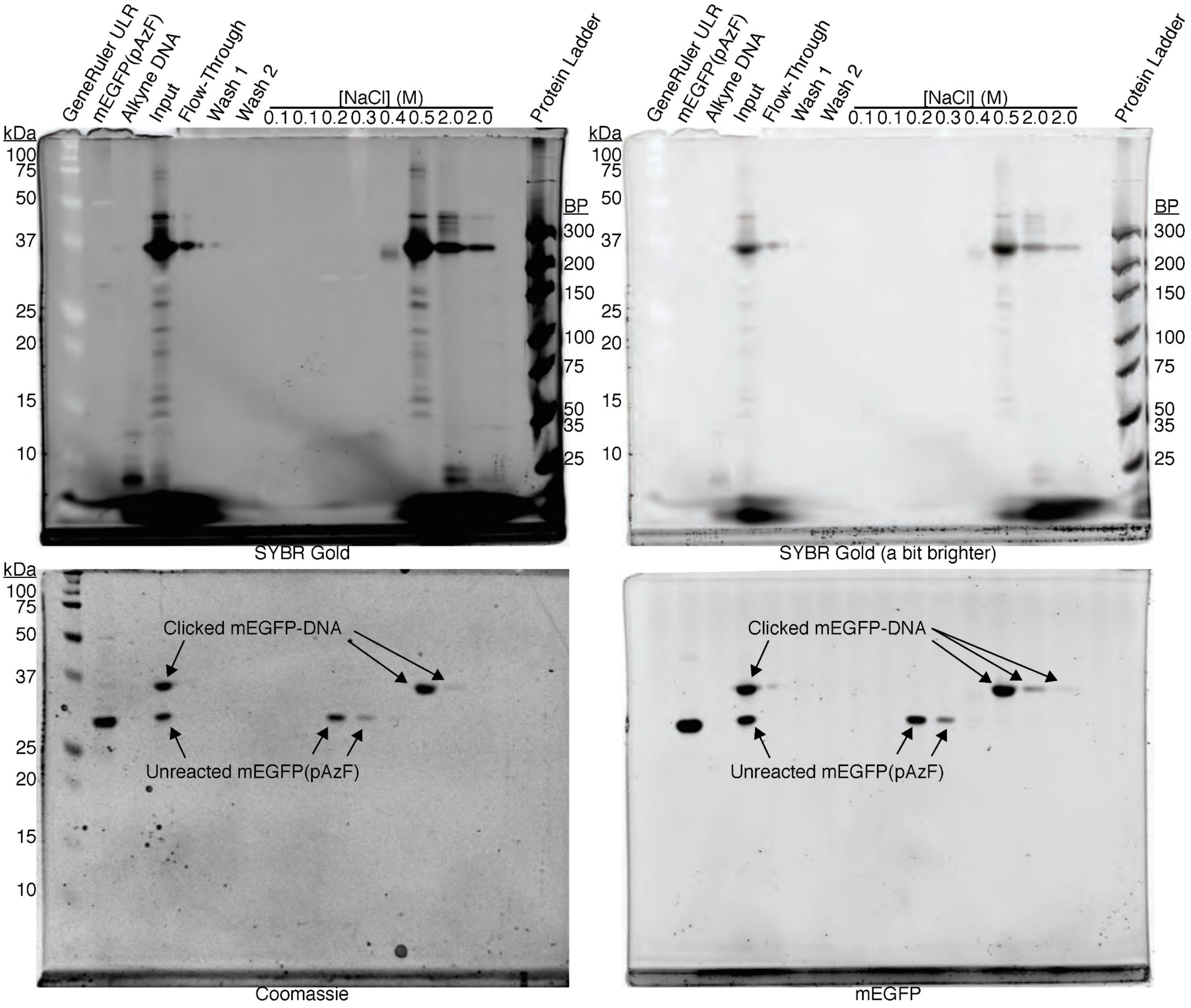

**Figure S6. mEGFP-DNA conjugates after anion-exchange purification.** The expected ~7 kDa shift in reacted mEGFP:anti-inner1 is clearly visible when comparing unreacted mEGFP(pAzF) and the Input reaction mixture. Unreacted mEGFP(pAzF) began to elute at an ionic strength of 0.2 M NaCl, while mEGFP:anti-Inner1 eluted above 0.4 M NaCl. Incomplete reaction of mEGFP(pAzF) is possibly due to imperfect incorporation of pAzF at the C-term and azide reduction *in vivo*.[16]

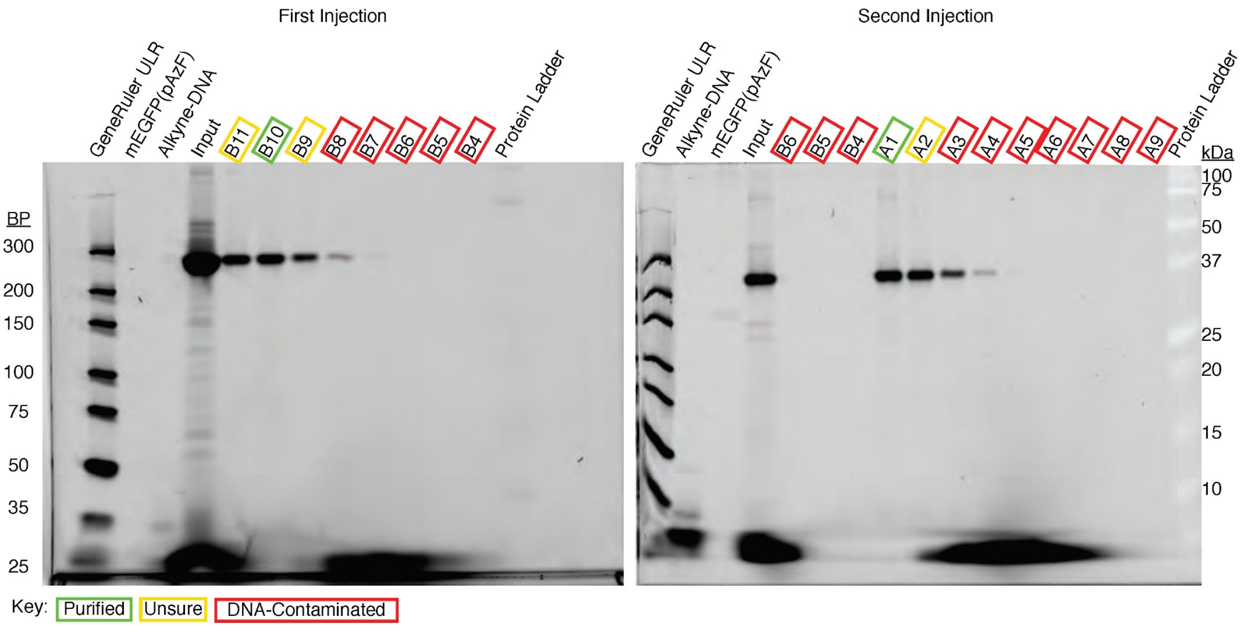

**Figure S7. mEGFP-DNA conjugates after size-exclusion purification.** Anion-exchanged reaction mixtures were purified (two rounds) by size-exclusion chromatography to remove unreacted alkyane-DNA. Gels were stained by Sybr Gold and imaged in the corresponding channel.

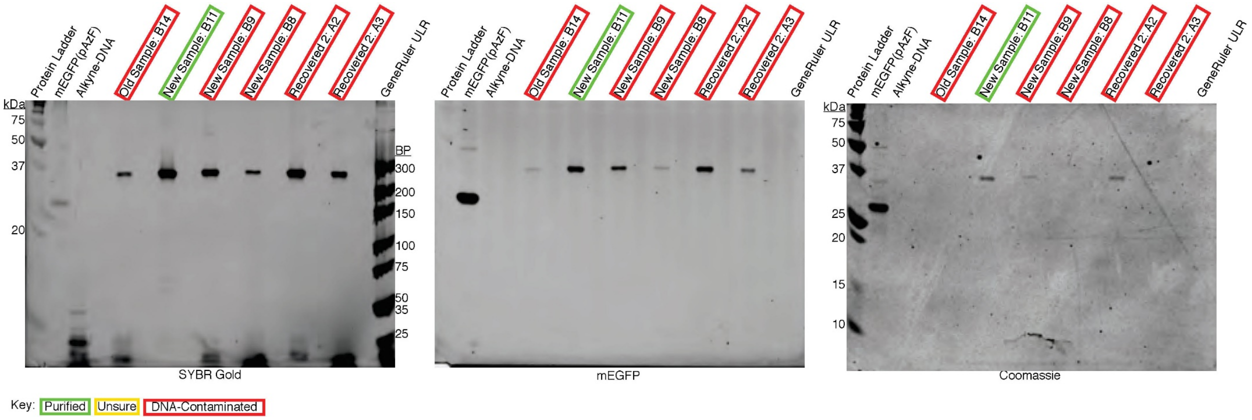

**Figure S8. Purified mEGFP-DNA conjugates.** Neither unreacted mEGFP nor DNA were found in purified fractions (B11).

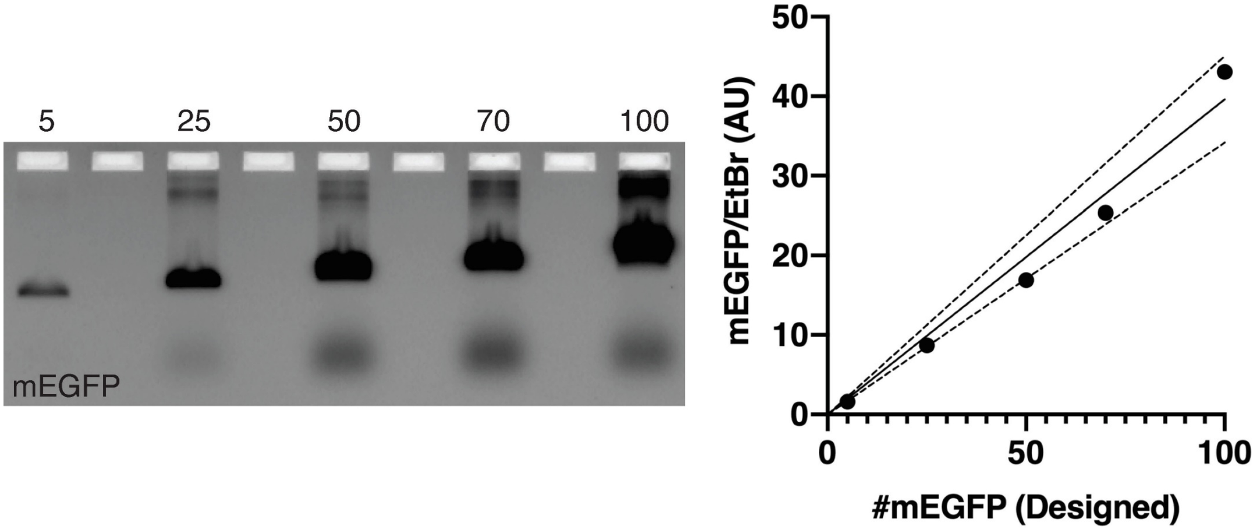

**Figure S9. mEGFP standards electrophoresed in an agarose gel.** DNA nanotubes designed to accomodate 5–100 mEGFP were labeled with mEGFP-antihandles and run in a non-denaturing agarose gel (left). After the mEGFP channel (488 nm) was imaged, the gel was stained with 0.5 μg/mL EtBr and imaged again (532 nm). The band intensities were quantified using ImageQuant TL, and normalized mEGFP fluorescence was plotted (right). A standard linear regression[11] yielded a slope of 0.3962 (solid line) with a 95% confidence interval (CI) of 0.3417 to 0.4508 (dashed lines).

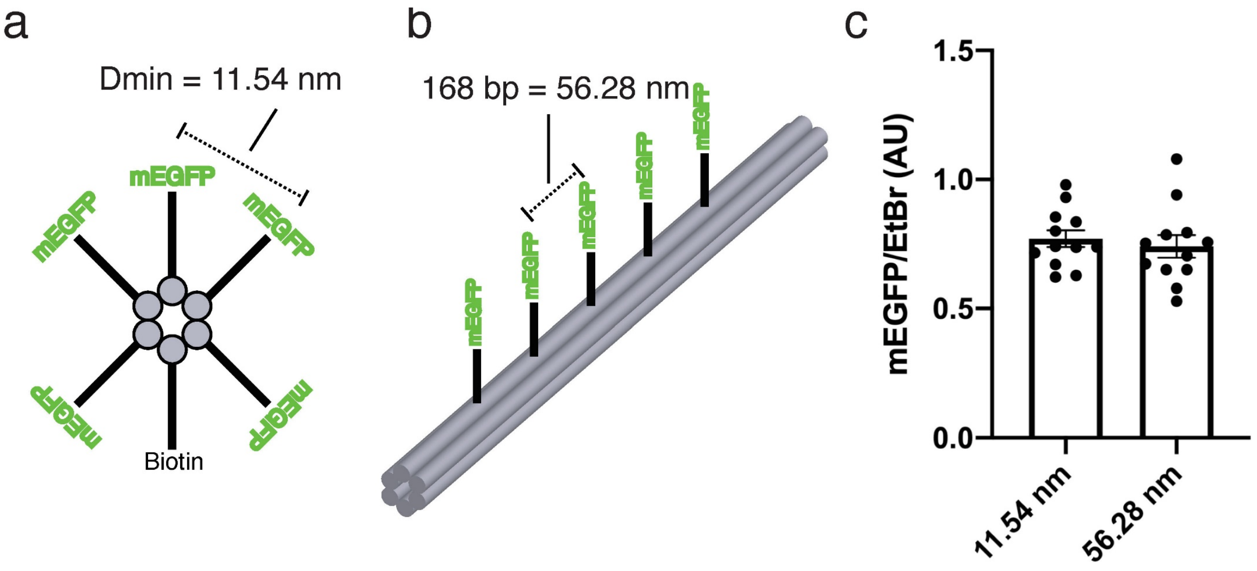

**Figure S10. Test for mEGFP self-quenching.** 6 Structures were designed to host 5× mEGFPs, 3 with the same inter-protein distances as the rest of this work (a), and 3 had mEGFP placed every ~56 nm (b). In each structure, unique handle locations were chosen to negate the influence of specific handle incorporation efficiencies. All 6 structures were run in quadruplicate in a non-denaturing agarose gel, and imaged before and after EtBr staining. Normalized mEGFP band intensities were nearly identical, regardless of spacing (mean±SEM), showing no sign of self-quenching.

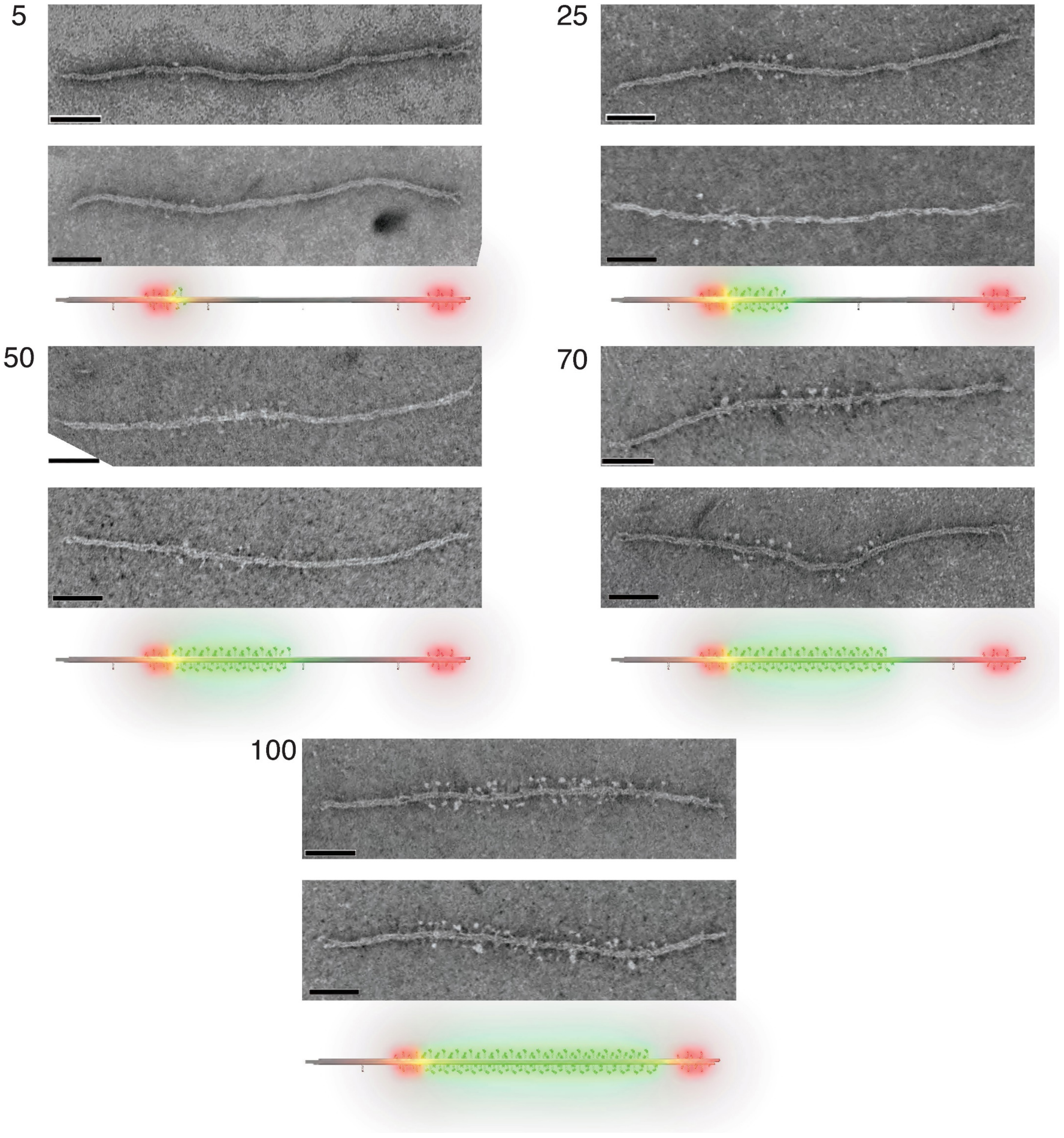

**Figure S11. TEM micrographs of mEGFP standards**. 5, 25, 50, 70, & 100 indicate the number of mEGFP each structure is designed to accommodate. Due to the resolution limit of negative-stain TEM, we do not expect to resolve every single mEGFP molecule. Nevertheless, in selected images, small dots (mEGFP) were found spanning the structures in the expected regions. Cartoon models were placed below micrographs (Green: GFP, Red: Alex Fluor 647) for direct comparison. Scale bars: 50 nm.

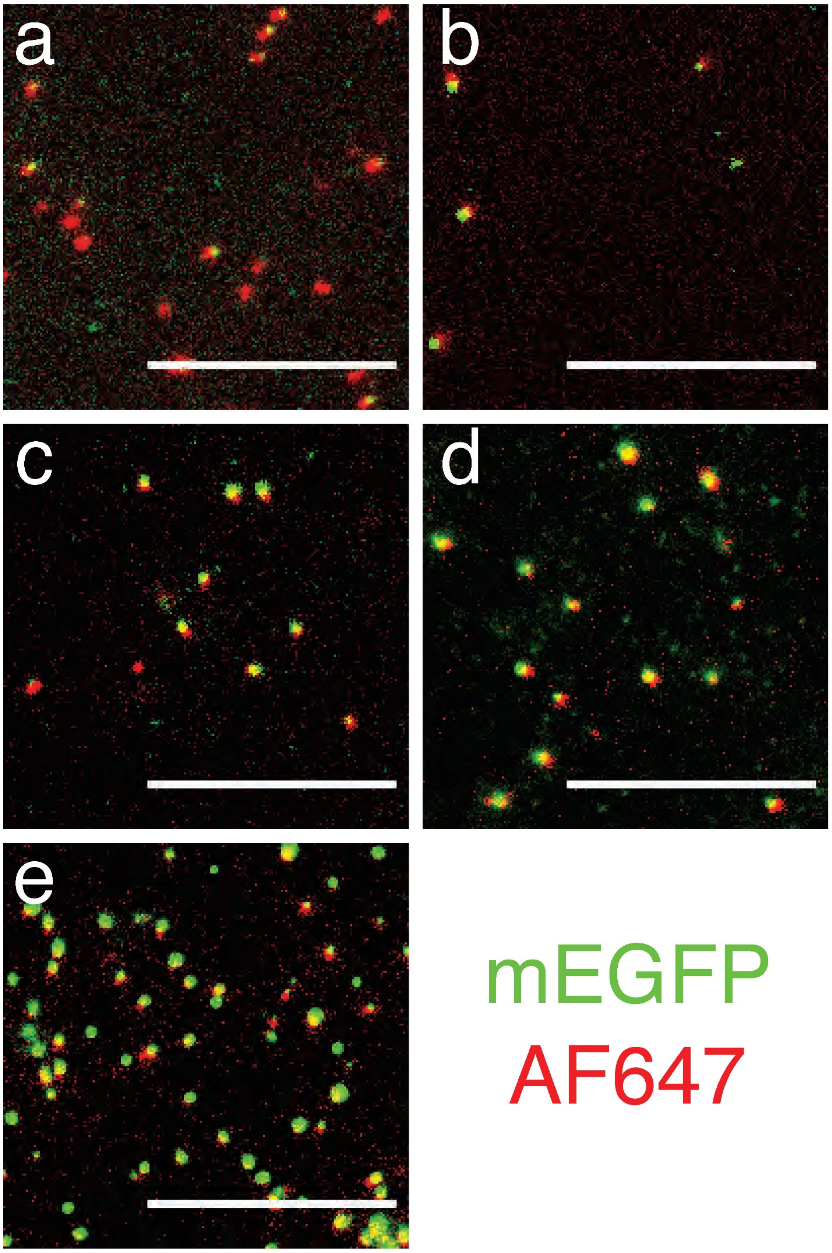

**Figure S12. Wide-field fluorescence micrographs of mEGFP standards.**. (a) 5×, (b) 25×, (c) 50×, (d) 70×, (e) 100×. Brightness and contrast for each image adjusted individually for clarity. Scale bars: 10 μm.

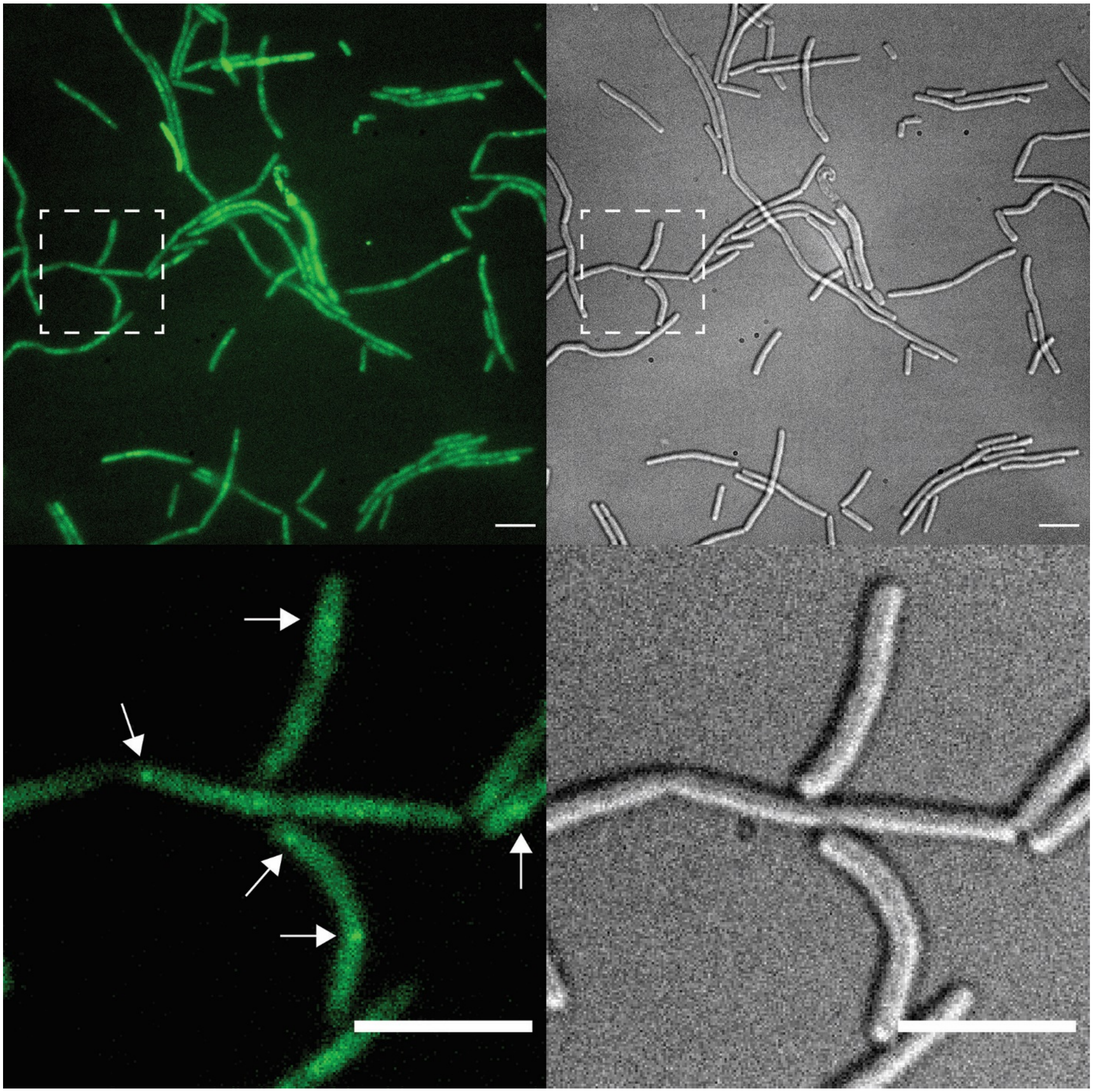

**Figure S13. Wide-field fluorescence micrographs of *B. subtilis***. Representative image of *B. subtilis* (strain NW001) cells expressing dnaC-mEGFP (puncta indicated by arrows). Left: mEGFP (488 nm); right: differential interference contrast. Images in the bottom row are magnified from areas within dotted rectangles. Brightness and contrast for each image adjusted individually for clarity. Unlike Figure 1d, the fluorescence brightness is rescaled not to highlight individual punctum, but to show the overall cell shapes. Scale bars: 10 μm.

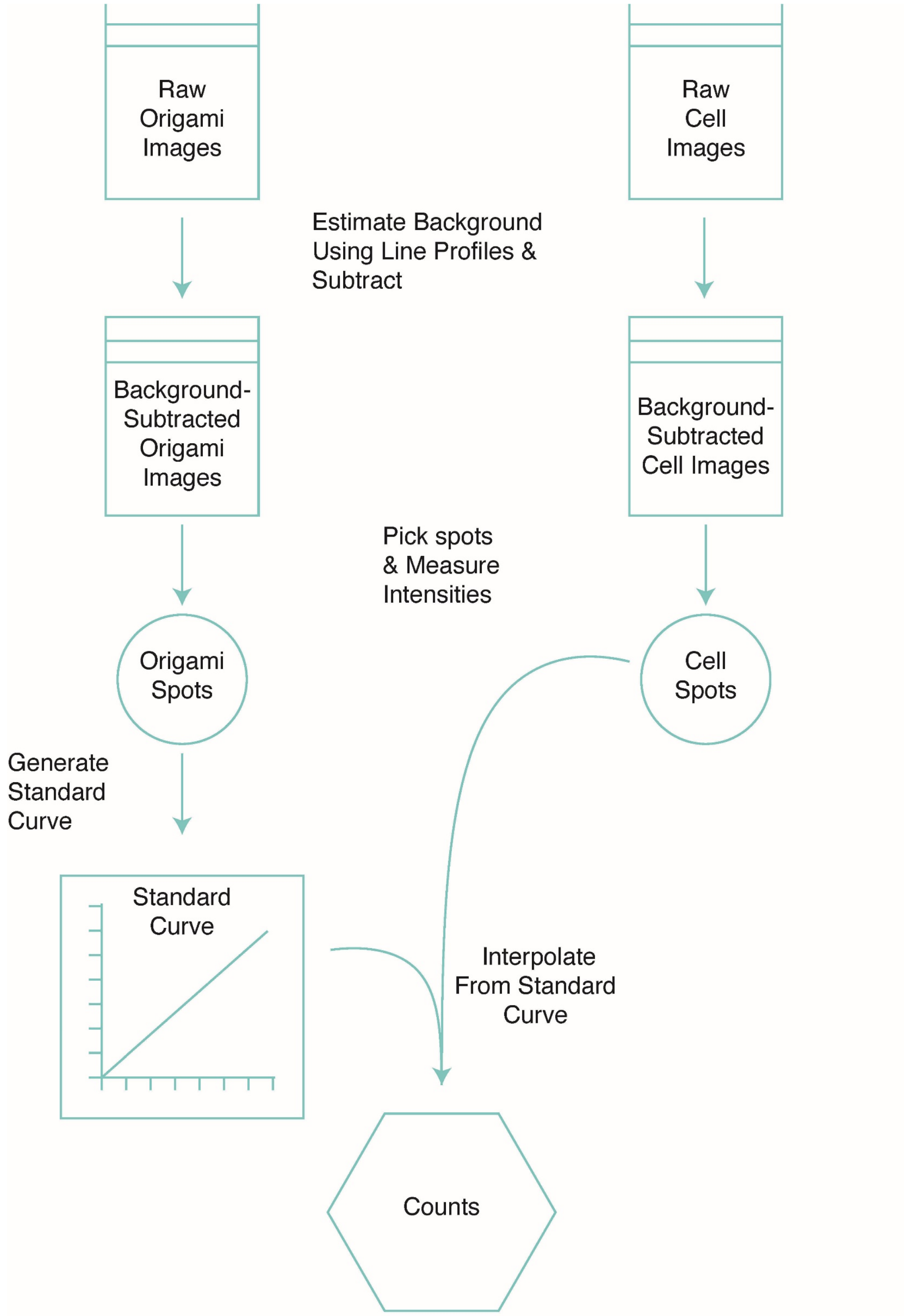

**Figure S14. Image processing pipeline for mEGFP labeling.** Background fluorescence from agar pads and cell auto-fluorescence was estimated by measuring line profiles spanning cells and DNA-origami standards, then subtracted from the image. Spots were picked using MicrobeJ[10], and were subsequently selected manually (see Materials and Methods for criteria). These intensities were fit to a sum of two Gaussians function using GraphPad Prism[11].

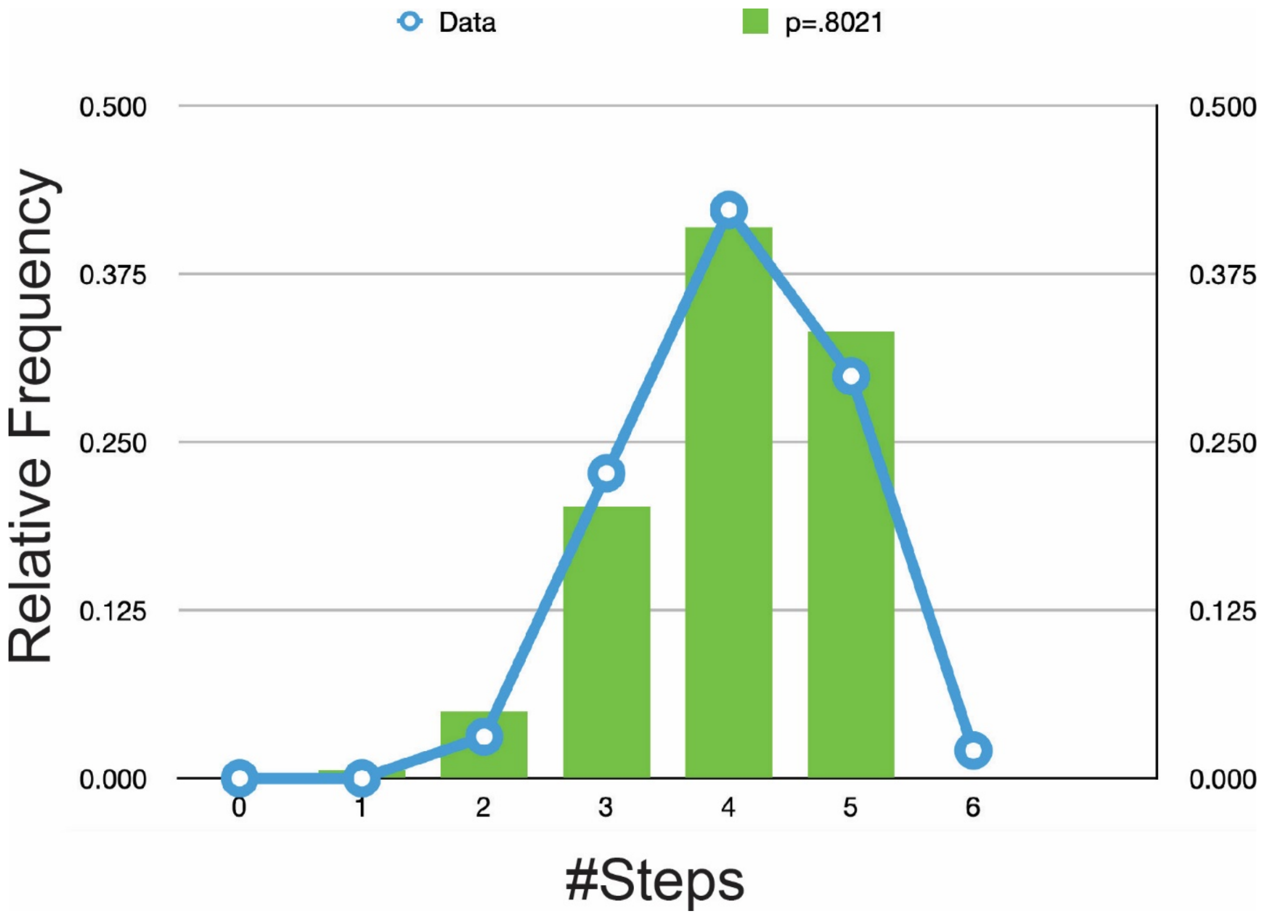

**Figure S15. Handle occupancy estimation.** Structures designed to host 5× mEGFP were labeled with Alexa Fluor 488-labeled antihandles and imaged on a Nikon TIRF microscope until fully bleached. The total photobleaching steps of the fluorescence traces were fit to a binomial function with MATLAB[17]. The best fit probability was p=0.8021, 95% CI = [0.7634 0.8370]. Note: 2 out of 97 traces showed 6 apparent steps, and were excluded from the binomial fit.

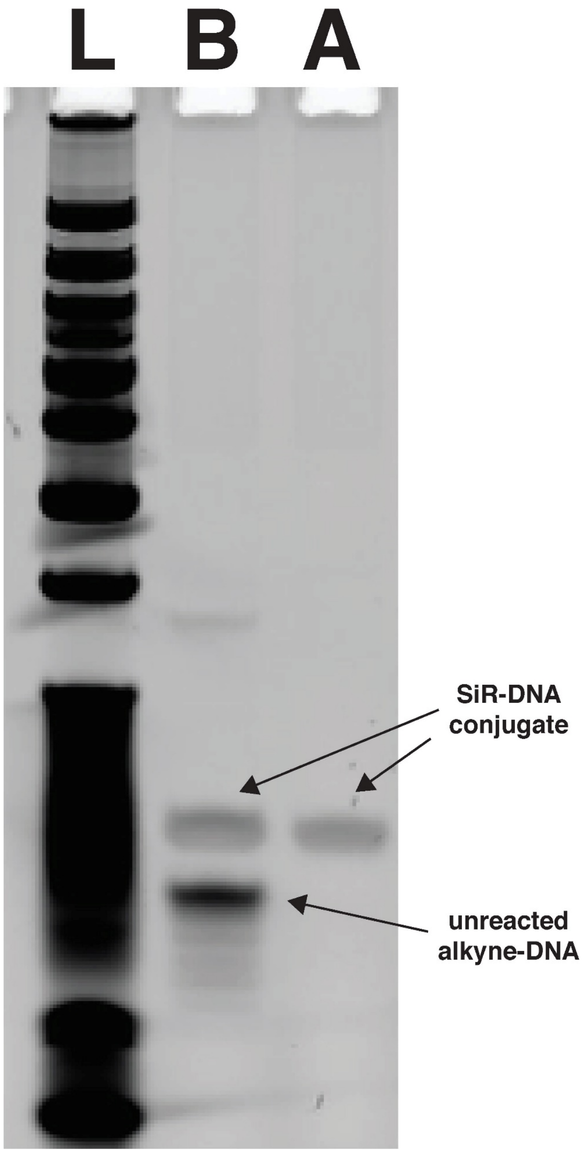

**Figure S16. SiR-DNA conjugate analyzed by PAGE.** SYBR Gold-stained 15% urea-PAGE gel of the SiR conjugate before (B) and after (A) purification. Ladder (L) is NEB Ultra Low Range DNA Ladder.

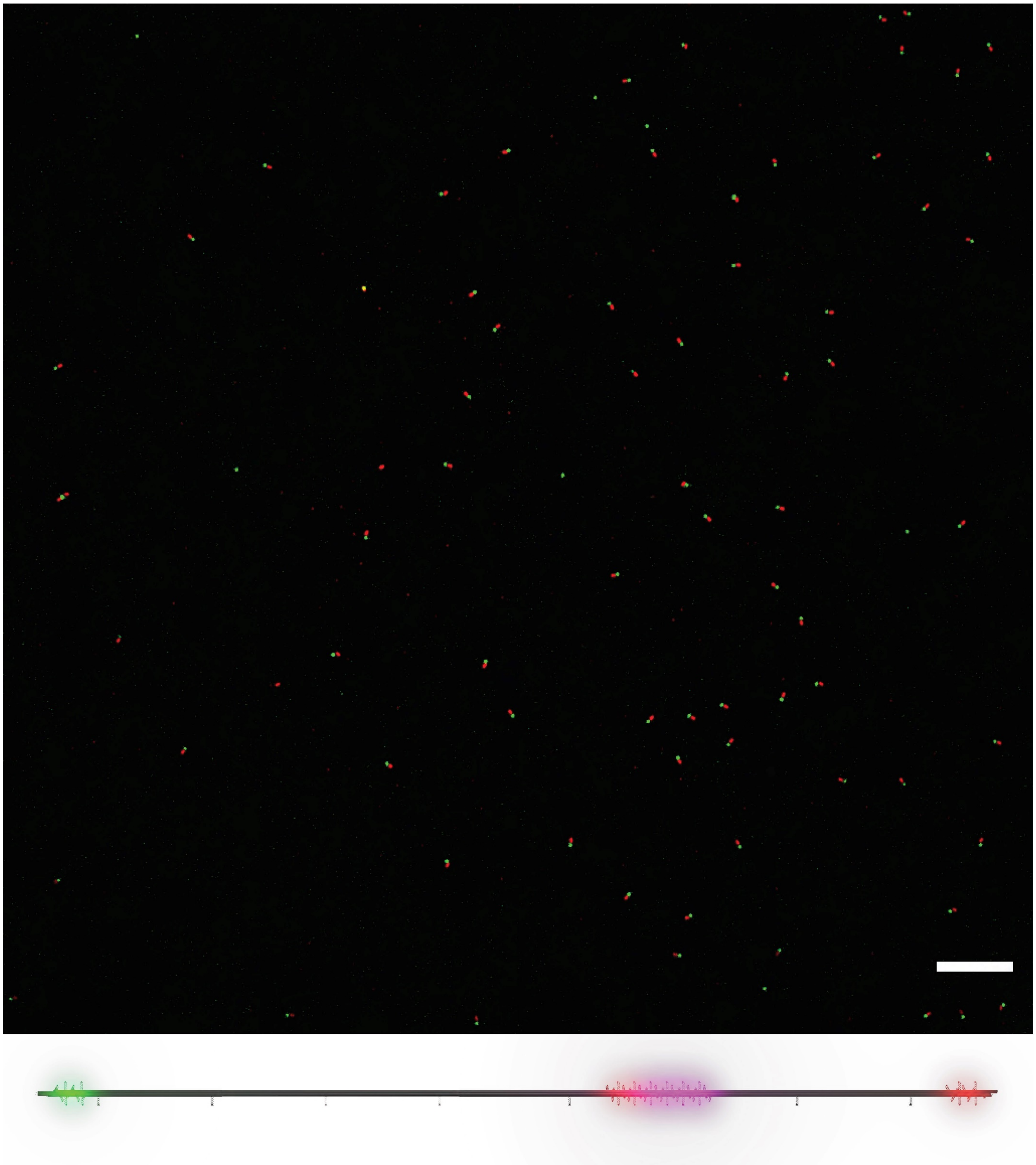

**Figure S17. Fluorescent barcode pattern on 25× SiR standard**. Top: confocal fluorescence micrograph showing the barcoding channels of Alexa Fluor 488 (green), and TAMRA (red). Scale bar: 10 μm. Bottom: A 3D model showing positions of barcoding fluorophores on the 25×SiR-labeled DNA-origami 6hb structure.

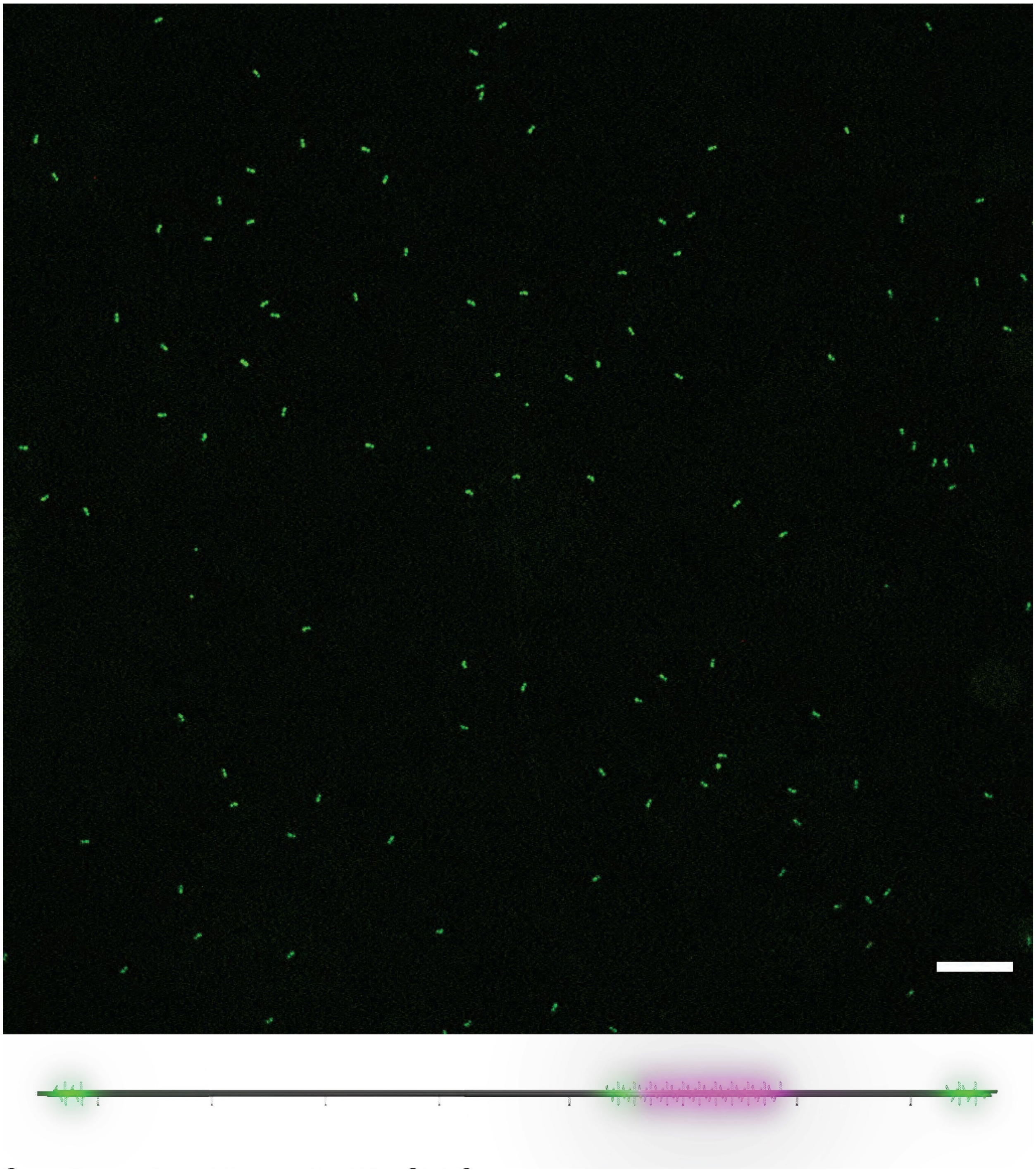

**Figure S18. Fluorescent barcode pattern on 50× SiR standard**. Top: confocal fluorescence micrograph showing the barcoding channel of Alexa Fluor 488 (green). Scale bar: 10 µm. Bottom: A 3D model showing positions of barcoding fluorophores on the 50×SiR-labeled DNA-origami 6hb structure.

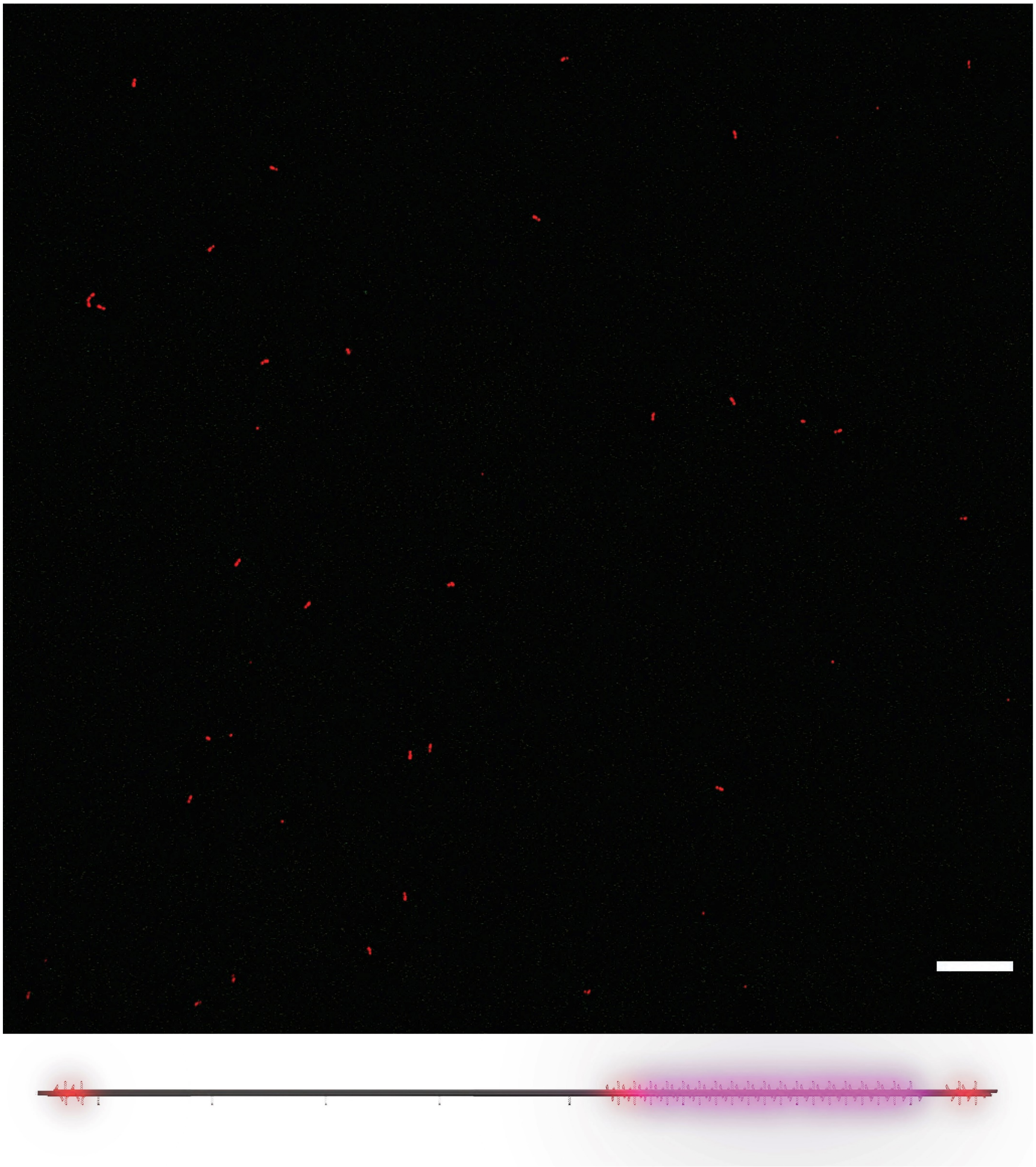

**Figure S19. Fluorescent barcode pattern on 100× SiR standard**. Top: confocal fluorescence micrograph showing the barcoding channel of TAMRA (red). Scale bar: 10 μm. Bottom: A 3D model showing positions of barcoding fluorophores on the 100×SiR-labeled DNA-origami 6hb structure.

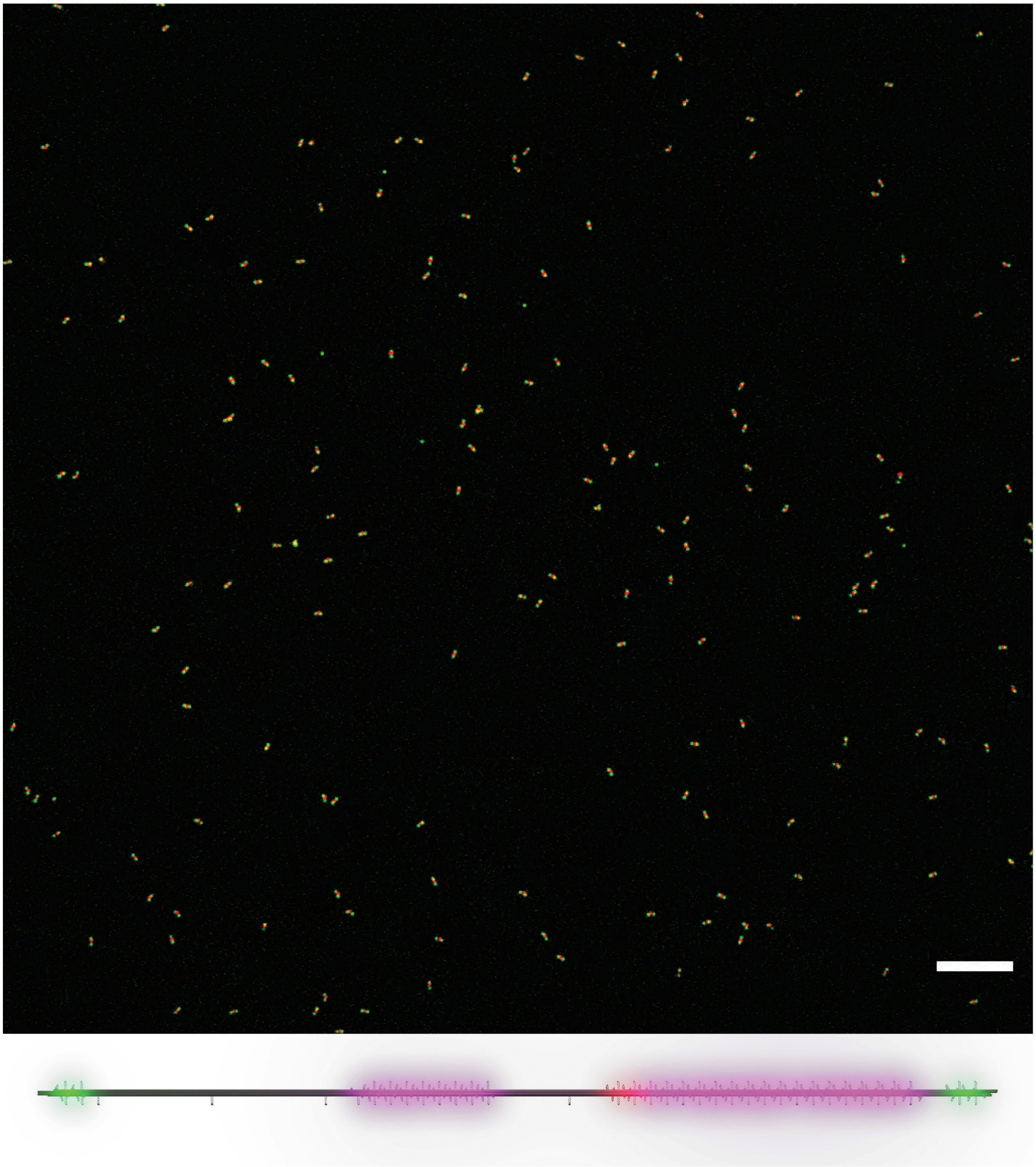

**Figure S20. Fluorescent barcode pattern on 150× SiR standard**. Top: confocal fluorescence micrograph showing the barcoding channels of Alexa Fluor 488 (green), and TAMRA (red). Scale bar: 10 μm. Bottom: A 3D model showing positions of barcoding fluorophores on the 150×SiR-labeled DNA-origami 6hb structure.

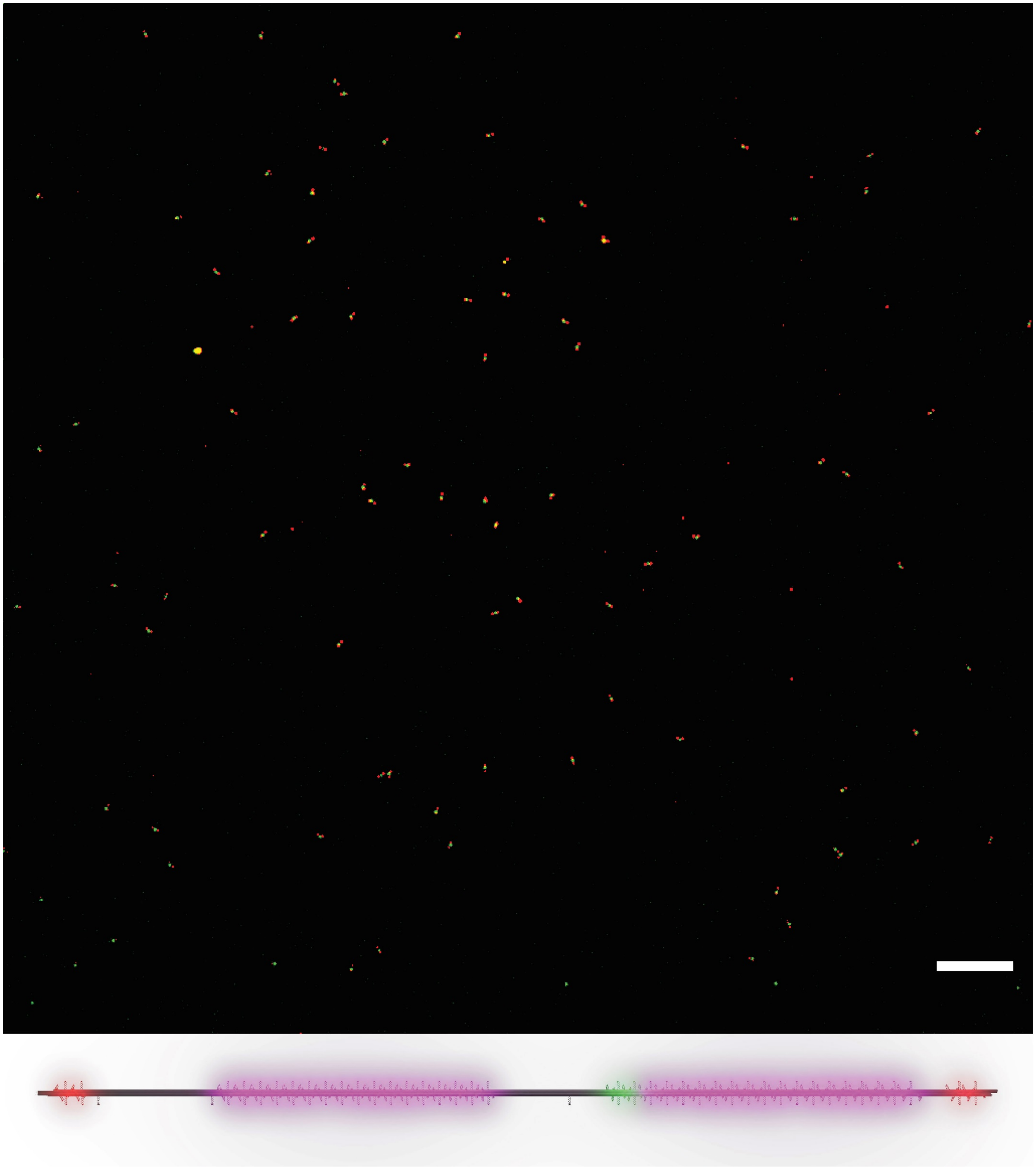

**Figure S21. Fluorescent barcode pattern on 200× SiR standard**. Top: confocal fluorescence micrograph showing the barcoding channels of Alexa Fluor 488 (green), and TAMRA (red). Scale bar: 10 μm. Bottom: A 3D model showing positions of barcoding fluorophores on the 200×SiR-labeled DNA-origami 6hb structure.

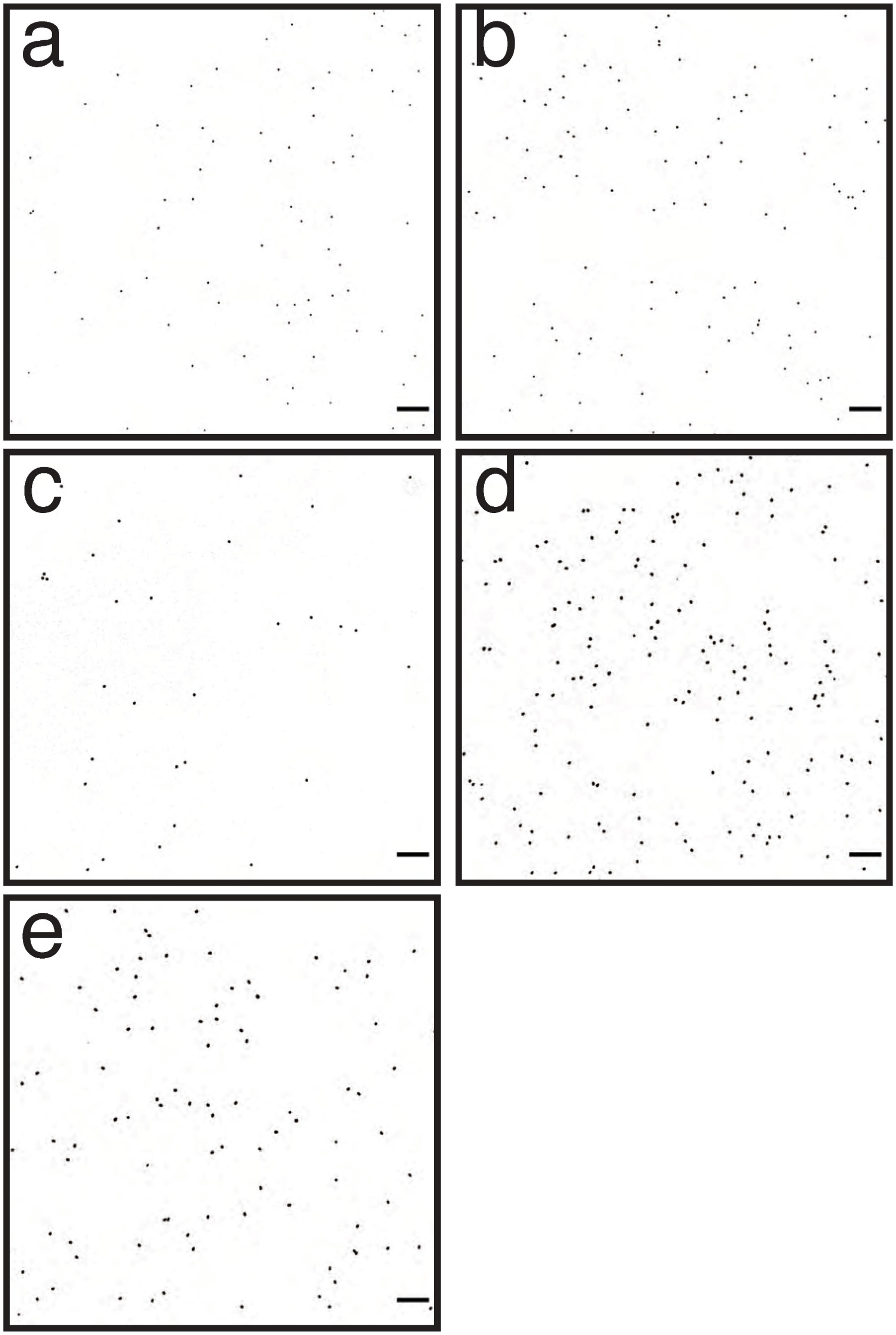

**Figure S22. Confocal fluorescence micrographs of SiR standards.** Panels show the SiR channel of Figure S17–S21, respectively: (a) 25× SiR, (b) 50× SiR, (c) 100× SiR, (d) 150× SiR, (e) 200× SiR. All panels adjusted to the same brightness and contrast levels. Scale bars: 10 µm.

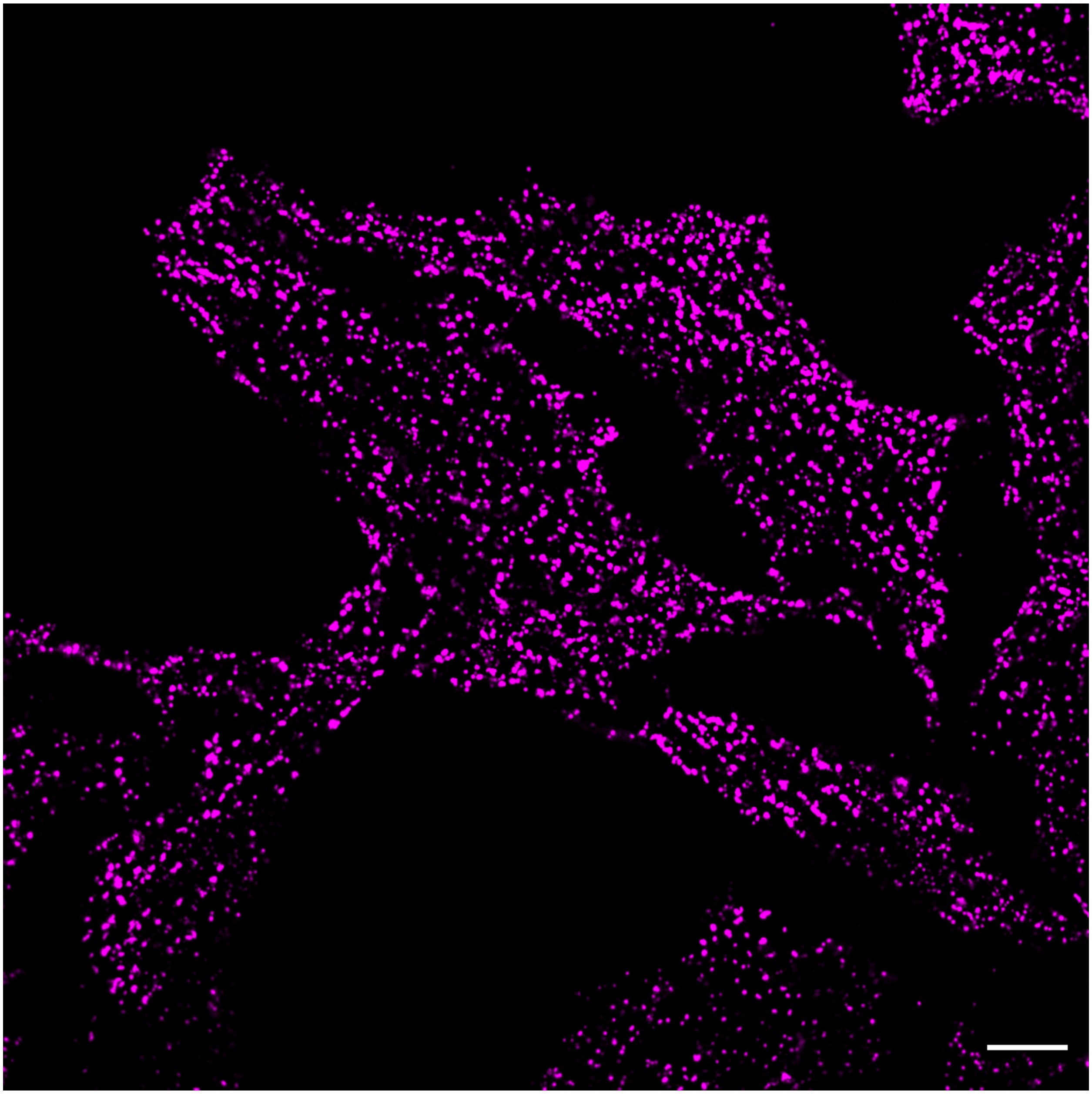

**Figure S23. A confocal fluorescence micrograph of Hela cells.** The focal plane is set close to the bottom of the dish. Scale bar: 10 μm.

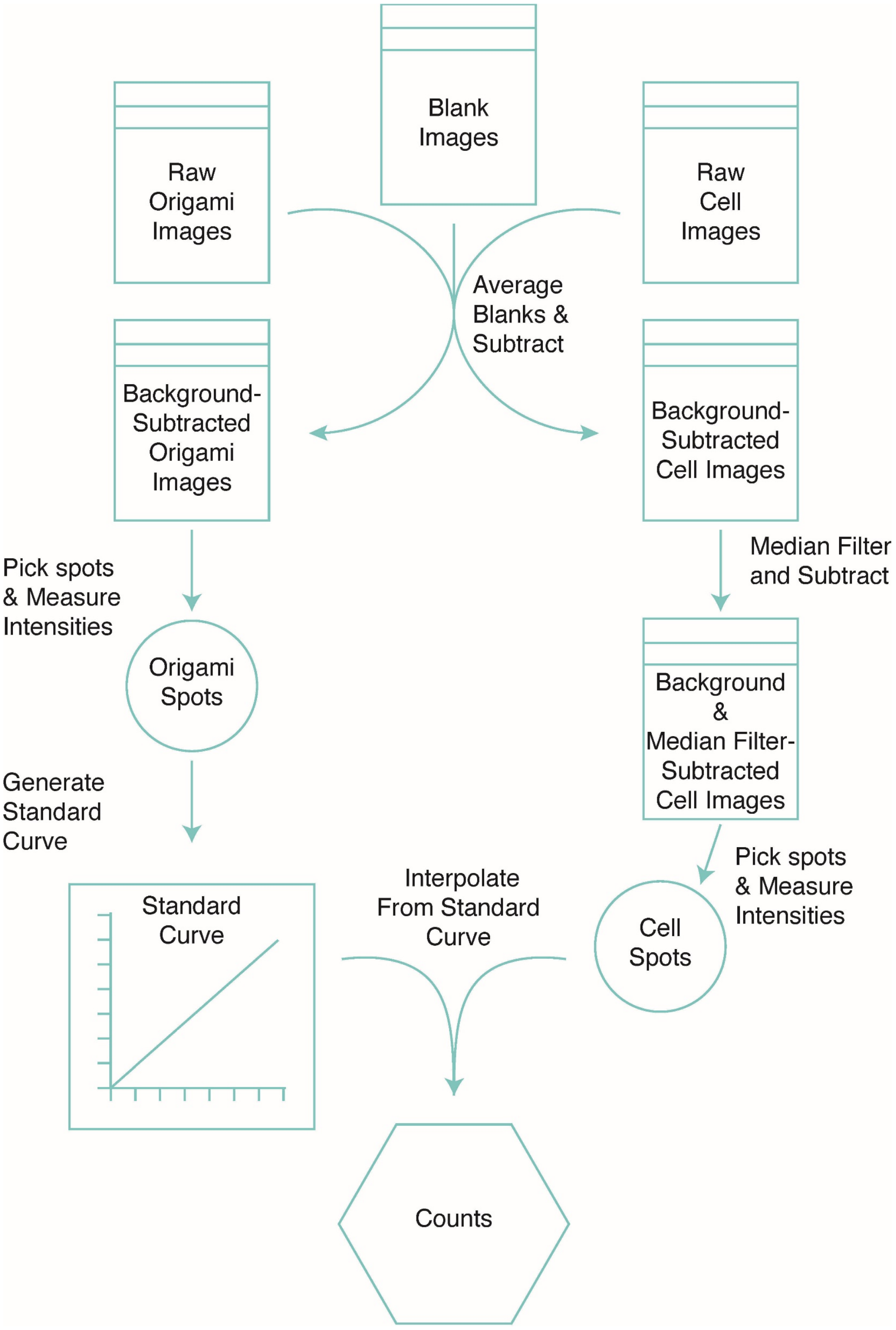

**Figure S24. Image processing pipeline for SiR labeling.** Background from media and the dish was approximated from empty dishes (blank images), and subtracted from all images containing SiR standards or HeLa cells. SiR puncta in DNA-origami images were picked using a custom TrackMate script, and used to construct the calibration curve. A background-subtracted image of cells was median filtered, and was subtracted from the cell images. CLC-SiR spots in cells were then picked manually. CLC-SiR intensities were then converted to molecule number using the calibration curve.

### Supplementary Tables

**Table S1: Handle and antihandle sequences**

| Name | Sequence (5'-3') |
|---|---|
| Outer 2 | **CTTCACACCACACTCCATCTA** |
| Inner 1 | **AAATTATCTACCACAACTCAC** |
| handle ix | **ACCTACTAACATAATCATCAC** |
| Non ATG 1 handle | **CGGTTGTACTGTGACCGATTC** |
| Anti Outer 2 | **TAGATGGAGTGTGGTGTGAAG** |
| Anti Inner 1 | **GTGAGTTGTGGTAGATAATTT** |
| anti-handle ix | **GTGATGATTATGTTAGTAGGT** |
| Non-ATG 1 antihandle | **GAATCGGTCACAGTACAACCG** |
| 5'Alkyne-anti Inner 1 | **/5Hexynyl/GTGAGTTGTGGTAGATAATTT** |
| 5' Biotin Non-ATG 1 | **/5Biosg/GAATCGGTCACAGTACAACCG** |
| 3'AF488-anti Inner 1 | **GTGAGTTGTGGTAGATAATTT/3AlexF488N/** |
| 5' AF488-anti handle ix | **/5Alex488N/TGTGATGATTATGTTAGTAGGT** |
| 5' TAMRA-anti handle ix | **/56-TAMN/GTGATGATTATGTTAGTAGGT** |
| 5' TAMRA-anti outer 2 | **/56-TAMN/TAGATGGAGTGTGGTGTGAAG** |
| 5' AF647-anti outer 2 | **/5Alex647N/TAGATGGAGTGTGGTGTGAAG** |

**Table S2: Linker DNA strands used in dimers**

| Name | Sequence (5'-3') |
|---|---|
| LD1.1 | CATTGCATGCCTGCGGAATTAGAGCCAGAAAGGTGAATTATC |
| LD1.2 | CCAGTGCCAAGCGATTTGAAATACCGACAGAAAAAGCCTGTT |
| LD1.3 | ACCGTCACCGACCCGAATCATAATTACTCGTGTGATAAATAA |
| LD1.4 | GGTTGAGCCATTTGAGGTCGACTCTAGATTGTAAAACGACGG |
| LD1.5 | GGCGTTAAATAACATCCCAGTCACGACGCCTTTGATAGCGAG |
| LD1.6 | AAGAATAAACACCGGCTTTTGCGGATCTGCAGGGAGTTAAA |

### Supplementary References

(1) Douglas, S. M.; Marblestone, A. H.; Teerapittayanon, S.; Vazquez, A.; Church, G. M.; Shih, W. M. Rapid Prototyping of 3D DNA-Origami Shapes with CaDNAno. *Nucleic acids research* **2009**, *37* (15), 5001–5006. https://doi.org/10.1093/nar/gkp436.

(2) Douglas, S. M.; Dietz, H.; Liedl, T.; Högberg, B.; Graf, F.; Shih, W. M. Self-Assembly of DNA into Nanoscale Three-Dimensional Shapes. *Nature* **2009**, *459* (7245), 414–418. https://doi.org/10.1038/nature08016.

(3) Stahl, E.; Martin, T. G.; Praetorius, F.; Dietz, H. Facile and Scalable Preparation of Pure and Dense DNA Origami Solutions. *Angewandte Chemie International Edition* **2014**, *53* (47), 12735–12740. https://doi.org/10.1002/anie.201405991.

(4) Lin, C.; Perrault, S. D.; Kwak, M.; Graf, F.; Shih, W. M. Purification of DNA-Origami Nanostructures by Rate-Zonal Centrifugation. *Nucleic acids research* **2013**, *41* (2), e40–e40. https://doi.org/10.1093/nar/gks1070.

(5) Lajoie, M. J.; Rovner, A. J.; Goodman, D. B.; Aerni, H.-R.; Haimovich, A. D.; Kuznetsov, G.; Mercer, J. A.; Wang, H. H.; Carr, P. A.; Mosberg, J. A.; Rohland, N.; Schultz, P. G.; Jacobson, J. M.; Rinehart, J.; Church, G. M.; Isaacs, F. J. Genomically Recoded Organisms Expand Biological Functions. *Science (New York, N.Y.)* **2013**, *342* (6156), 357–360. https://doi.org/10.1126/science.1241459.

(6) Amiram, M.; Haimovich, A. D.; Fan, C.; Wang, Y.-S.; Aerni, H.-R.; Ntai, I.; Moonan, D. W.; Ma, N. J.; Rovner, A. J.; Hong, S. H.; Kelleher, N. L.; Goodman, A. L.; Jewett, M. C.; Söll, D.; Rinehart, J.; Isaacs, F. J. Evolution of Translation Machinery in Recoded Bacteria Enables Multi-Site Incorporation of Nonstandard Amino Acids. *Nature biotechnology* **2015**, *33* (12), 1272–1279. https://doi.org/10.1038/nbt.3372.

(7) Presolski, S. I.; Hong, V. P.; Finn, M. G. Copper-Catalyzed Azide-Alkyne Click Chemistry for Bioconjugation. *Current protocols in chemical biology* **2011**, *3* (4), 153–162. https://doi.org/10.1002/9780470559277.ch110148.

(8) Lin, C.; Jungmann, R.; Leifer, A. M.; Li, C.; Levner, D.; Church, G. M.; Shih, W. M.; Yin, P. Submicrometre Geometrically Encoded Fluorescent Barcodes Self-Assembled from DNA. *Nat Chem* **2012**, *4* (10), 832–839. https://doi.org/10.1038/nchem.1451.

(9) Mangiameli, S. M.; Veit, B.T.; Merrikh, H.; Wiggins, P. A. The Replisomes remain spatially proximal throughout the Cell Cycle in Bacteria. *PLOS Genetics*. **2017**, *13*, e1006582. https://doi.org/10.1371/journal.pgen.1006582.

(10) Ducret, A.; Quardokus, E. M.; Brun, Y. V. MicrobeJ, a Tool for High Throughput Bacterial Cell Detection and Quantitative Analysis. *Nat. Microbiol.* **2016**, 1–7. https://doi.org/10.1038/nmicrobiol.2016.77.

(11) *GraphPad Prism* version 8.4.3 for macOS; GraphPad Software: San Diego, CA USA, 2020.

(12) Cong, L.; Ran, F. A.; Cox, D.; Lin, S.; Barretto, R.; Habib, N.; Hsu, P. D.; Wu, X.; Jiang, W.; Marraffini, L. A.; Zhang, F. Multiplex Genome Engineering Using CRISPR/Cas Systems. *Science (New York, N.Y.)* **2013**, *339* (6121), 819–823. https://doi.org/10.1126/science.1231143.

(13) Takakura, H.; Zhang, Y.; Erdmann, R. S.; Thompson, A. D.; Lin, Y.; McNellis, B.; Rivera-Molina, F.; Uno, S.; Kamiya, M.; Urano, Y.; Rothman, J. E.; Bewersdorf, J.; Schepartz, A.; Toomre, D. Long Time-Lapse Nanoscopy with Spontaneously Blinking Membrane Probes. *Nature Biotechnol.* **2017**, 773–780. https://doi.org/10.1038/nbt.3876.

(14) Saffarian, S.; Cocucci, E.; Kirchhausen, T. Distinct Dynamics of Endocytic Clathrin-Coated Pits and Coated Plaques. *PLOS Biology* **2009**, *7* (9), e1000191-18. https://doi.org/10.1371/journal.pbio.1000191.

(15) Gan, L.; Chao, T.-C.; Camacho-Alanis, F.; Ros, A. Six-Helix Bundle and Triangle DNA Origami Insulator-Based Dielectrophoresis. *Analytical Chemistry* **2013**, *85* (23), 11427–11434. https://doi.org/10.1021/ac402493u.

(16) Milles, S.; Tyagi, S.; Banterle, N.; Koehler, C.; VanDelinder, V.; Plass, T.; Neal, A. P.; Lemke, E. A. Click Strategies for Single-Molecule Protein Fluorescence. *J. Am. Chem. Soc.* **2012**, *134* (11), 5187–5195. https://doi.org/10.1021/ja210587q.

(17) *The MathWorks MATLAB* version 9.8.0.1359463 (R2020a) Update 1 for macOS; The MathWorks, Inc.: Natick, MA USA, 2020.
